# PDGFRA Defines the Mesenchymal Stem Cell Kaposi’s Sarcoma Progenitors by Enabling KSHV Oncogenesis in an Angiogenic Environment

**DOI:** 10.1101/789826

**Authors:** Julian Naipauer, Santas Rosario, Sachin Gupta, Courtney Premer, Omayra Mendez, Mariana Schlesinger, Virginia Ponzinibbio, Vaibhav Jain, Lauren Gay, Rolf Renne, Ho Lam Chan, Lluis Morey, Daria Salyakina, Martin Abba, Sion Williams, Joshua M. Hare, Pascal J Goldschmidt-Clermont, Enrique A. Mesri

**Affiliations:** Tumor Biology Program, Sylvester Comprehensive Cancer Center and Miami Center for AIDS Research, Department of Microbiology and Immunology, Miami, FL 33136; UM-CFAR/ Sylvester CCC Argentina Consortium for Research and Training in Virally induced AIDS-Malignancies, Miami, FL 33136; Interdisciplinary Stem Cell Institute; University of Miami Miller School of Medicine, Miami, FL 33136; Department of Molecular Genetics and Microbiology, University of Florida, Gainesville, FL 32610, USA.; Department of Human Genetics, Miami, FL 33136; Neurology Basic Science Division, Sylvester Comprehensive Cancer Center; University of Miami Miller School of Medicine, Miami, FL 33136; Centro de Investigaciones Inmunológicas Básicas y Aplicadas, Facultad de Ciencias Médicas, Universidad Nacional de La Plata, La Plata, Argentina.

## Abstract

Kaposi’s sarcoma (KS) is an AIDS-defining cancer caused by the KS-associated herpesvirus (KSHV). Unanswered questions regarding KS are its cellular ontology and the conditions conducive to viral oncogenesis. We identify PDGFRA(+)/SCA-1(+) bone marrow-derived mesenchymal stem cells (Pα(+)S MSCs) as KS spindle-cell progenitors and found that pro-angiogenic environmental conditions typical of KS are critical for KSHV sarcomagenesis. This is because growth in KS-like conditions generates a de-repressed KSHV epigenome allowing oncogenic KSHV gene expression in infected Pα(+)S MSCs. Furthermore, these growth conditions allow KSHV-infected Pα(+)S MSCs to overcome KSHV-driven oncogene-induced senescence and cell cycle arrest via a PDGFRA-signaling mechanism; thus identifying PDGFRA not only as a phenotypic determinant for KS-progenitors but also as a critical enabler for viral oncogenesis.

**AUTHOR SUMMARY:** Identification of the KS progenitor cell creates the possibility of studying viral oncogenesis and its determinants from its initial steps as a continuum. It also increases our understanding of pathogenic mechanisms and disease preferential tropism. Hereby we identify Pα(+)S-MSCs as KS progenitors, in which KSHV infection has oncogenic consequences; only when these cells are in a pro-angiogenic environment in which PDGFRA activation enables an oncogenic de-repressed KSHV epigenome. These results identify a KS-progenitor population in the Pα(+)S-MSCs and point to pro-angiogenic environmental conditions as essential for oncogenic viral gene expression and transformation. We designed a novel model of KSHV oncogenesis, creating a very robust platform to identify KSHV oncogenic pathways and their relationship with cellular lineages and extracellular growth environments.

## INTRODUCTION

Viral cancers account for up to 12% of all human cancers and are characterized by the long incubation periods and the fact that the majority of infected individuals do not develop cancer. This is consequence of the need for specific host environmental factors or conditions such as immunosuppression, which are necessary to enable the expression of the oncogenic viral gene expression programs leading to full viral-mediated cellular transformation [1]. Kaposi’s sarcoma (KS) is an AIDS-defining cancer and a major global health challenge caused by the Kaposi’s sarcoma-associated herpesvirus (KSHV) [2–4]. It is characterized by the proliferation of spindle-shaped cells (SC), inflammatory infiltrate and abundant angiogenesis with blood vessel erythrocyte extravasation [2–5]. KS presents in 4 different clinical forms: classical, endemic, iatrogenic and epidemic/AIDS-associated. Classical KS affects mostly elderly individuals of Ashkenazy Jews or Mediterranean descent and more recently at-risk populations such as men who have sex with men (MSM). Endemic KS affects children, men, and women in Sub-Saharan Africa. Iatrogenic KS is characteristic of transplant immunosuppression, in particular, renal transplant, and epidemic or AIDS-associated KS predominantly affects MSMs infected with HIV [4]. AIDS-associated immunosuppression and HIV constitute important KS co-factors, yet other host factors may account for the oncogenicity of KSHV and HIV co-infection in specific “at-risk” populations [6], Although the incidence of AIDS-KS in the western world has declined since the implementation of ART, more than 50% of advanced AIDS-KS patients never achieve total remission [6–8], Moreover, KSHV prevalence and KS appear to be increasing in ART-treated HIV-infected patients with controlled viremias [9, 10]. Critical pending questions on KS are its cellular ontology and the 4 conditions conducive to viral pathogenicity, which are important to understanding KSHV oncogeneic mechanisms that could lead to prevention approaches or the discovery of therapeutic targets.

The origins of KS spindle cells (SC) have long been debated, as these cells express markers of both lymphatic and blood vessel endothelium (podoplanin, VEGFR3, VEGF C and D, CXCR4, DLL4, VEGFR1, CXCL12, CD34)[11, 12], as well of dendritic cells (Factor XIII), macrophages (CD68), smooth muscle cells (SMA)[2] and mesenchymal stem cells (vimentin, PDGFRA)[13, 14]. This remarkable heterogeneity, together with the multifocal manifestation of many KS cases, suggests the existence of a circulating progenitor such as mesenchymal stem cells or endothelial cell progenitors [6, 15–17]. Spindle cell precursors were found to be increased in the blood of AIDS-KS patients, which upon KSHV infection and or inflammatory conditions may further differentiate into endothelial, smooth muscle, fibroblastic and myofibroblastic cells [18–20].

KSHV encodes a plethora of latent and lytic genes with pathogenic and oncogenic potential [2, 3]. KS lesions are composed of SC latently infected with KSHV, as well as cells expressing lytic genes that have been implicated in the development of the KS angioproliferative phenotype via paracrine and autocrine mechanisms [2, 3, 5, 21–23]. These mechanisms are mediated in part by the ability of lytic viral genes such as the G protein-coupled receptor (VGPCR/ORF74), K1 and K15, to upregulate angiogenesis and KS-cell growth factors [2, 3, 14, 21]. Although KSHV infection results in important morphological and transcriptional changes that convey traits of malignant transformation, few KSHV-infected cellular types had become fully tumorigenic [2, 5]. They are the basis for models of KSHV-5 tumorigenesis in murine, rat and human cells [24–28]. In a KSHV tumorigenesis model in nude mice generated by transfecting KSHVBac36 to mouse endothelial lineage cells [26], we found that malignancy was only manifested *in vivo* and occurred with concomitant upregulation of oncogenic KSHV lytic genes, angiogenesis growth factors and their tyrosine kinase receptors that are characteristic of human KS lesions [2, 26]. Using this model we show that the most prominently activated tyrosine kinase was PDGFRA [14], which was activated by lytic KSHV genes via a ligand-mediated mechanism and necessary for KSHV sarcomagenesis [14]. More importantly, PDGFRA was prominently expressed and phosphorylated in the vast majority of AIDS-KS tumors [14]. This together with the relative success of Imatinib Phase II trials targeting this receptor in KS [29]and the fact that PDGFRA is a driver of many sarcomas identified PDGFRA as an oncogenic driver in KS [14].

Viral cancers like KS are the consequence of infection with oncoviruses that evolved powerful mechanisms to persist and replicate through deregulation of host oncogenic pathways-such as PDGFRA signaling-conveying cancer hallmarks to the infected cell [1]. In searching for the identity of the oncogenic KS progenitor, we reasoned that if KSHV evolved its molecular machinery to activate PDGFRA signaling, a plausible oncogenic progenitor should be a pluripotent cell where PDGFRA plays a major proliferative and survival role, such as mesenchymal stem cell (MSC) [30]. This would also be consistent with the mesenchymal origin of sarcomas, many of which are driven by PDGFRA [31–33]. PDGFRA is a specific mesenchymal lineage phenotypic marker, and together with the stem cell antigen-1 (Sca-1), is used to identify pluripotent mesenchymal stem cells (Pα(+)S-MSCSX30]. Pα(+)S-MSCs have primitive characteristics consistent with conventional MSC populations and an *in vitro* differentiation 6 assay showed that single Pα(+)S-derived MSCs differentiated not only into chondrocytes and osteocytes but also into endothelial cells [30]. We hypothesized that PDGFRA-positive/SCA1-positive MSCs would likely serve as a natural target of KSHV infection, thus prompting oncogenic transformation via a PDGFRA-driven mechanism. The possibility of a mesenchymal progenitor for KS has been suggested by the work of several groups [28, 34–38]. In particular, mesenchymal and precursor markers are parts of the immunohistochemical features of KS [34]. In addition, rat and human MSCs from bone marrow and other origins are susceptible to KSHV infection [28, 35, 37, 38] and KSHV infection promoted multi-lineage differentiation and mesenchymal-to-endothelial transition into human oral MSC, providing evidence for human oral MSCs being a potential origin of Kaposi sarcoma cells [36].

Hereby we study the infection of murine and human PDGFRA+ MSC progenitors to generate a *“de novo”,* cell type-defined tumorigenesis system that could have the advantage to follow the process of tumorigenesis from a primary non-transformed cell, as well as identifying oncogenesis environmental conditions and molecular mechanisms critical for this process. In the present manuscript, we identify PDGFRA(+)/SCA-1(+) bone marrow-derived mesenchymal stem cells (Pα(+)S MSCs) as KS spindle cell progenitors and we also identify pro-angiogenic environmental conditions typical of KS as critical for KSHV sarcomagenesis. We found that growth in KS-like conditions generates a de-repressed KSHV epigenome, allowing oncogenic KSHV gene expression in infected Pα(+)S MSCs. Furthermore, these growth conditions allow KSHV-infected Pα(+)S MSCs to overcome KSHV-driven oncogene-induced senescence and cell cycle arrest via a PDGFRA-signaling mechanism; thus identifying PDGFRA not only as a phenotypic determinant for KS-progenitors but also as a critical enabler for viral oncogenesis.

## RESULTS

### KSHV latency establishment in mouse bone marrow-derived MSC

In order to identify spindle-cell KS progenitors, we purified adherent spindle-shaped mouse bone marrow-derived MSCs, from different mouse donors, that were positive for PDGFRA (60%) and the stem cell antigen SCA-1 (100%) (Figure 1A), as previously described [30, 39]· We sorted for PDGFRA-positive (Pα(+)S) and PDGFRA-negative (Pα(-)S) MSCs populations (Figure 1A), after infection with rKSHV.219 [40] the sorted populations were cultured in mesenchymal stem cell maintenance media (MSC-media). We monitored KSHV infection by examining the expression of a GFP cassette in the viral genome, and KSHV lytic infection through expression of RFP under the lytic viral PAN promoter. 72 hours post infection (hpi), we added puromycin to select for infected cells. Using this method, we successfully generated mouse bone marrow-derived MSC latently infected with KSHV (K-Pα(-)S and K-Pα (+)S) (Figure 1B). After several passages, these latently infected cultures continued to express GFP and the KSHV protein LANA, indicating the establishment of latent KSHV persistent infection (Figure 1C). Importantly, we did not detect RFP expression in either infected cell lines indicating stable KSHV latency. To study KSHV oncogenesis in these cells, we performed tumorigenic analysis and found that KSHV-uninfected and KSHV-infected cell populations that were PDGFRA-positive or PDGFRA-negative did not form tumors when subcutaneously injected in nude mice (S1 Figure).

**Figure 1.**
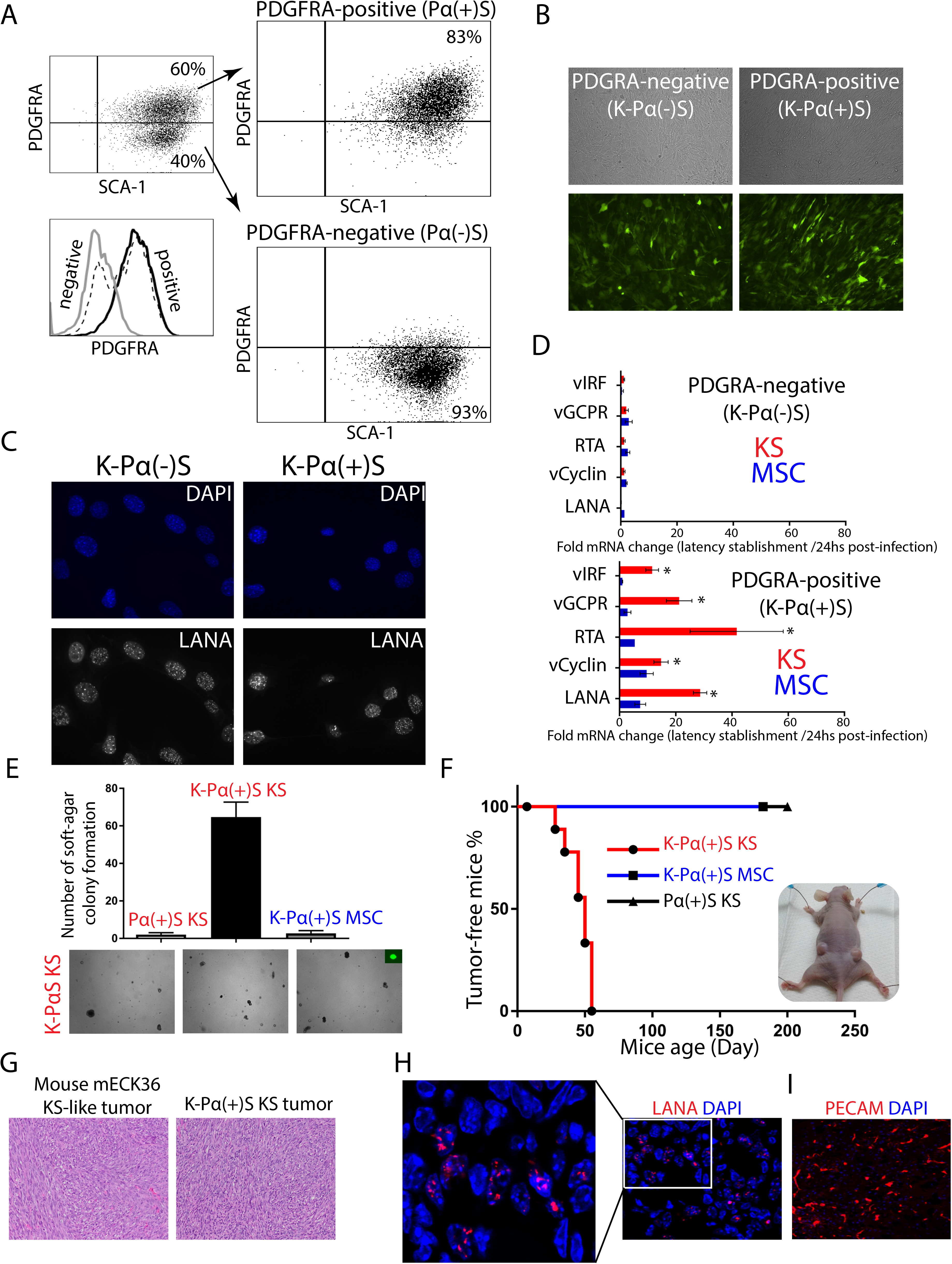
KSHV infection is only tumorigenic in infected MSC PDGFRA-positive lineagegrown in pro-angiogenic KS-like environment. A) Mouse bone marrow-derived mesenchymal stem cells were stained for PDGFRA (Pα) and SCA-1 (S) expression. Mouse MSC Sca-1-positive, PDGFRA-positive (Pα(+)S) and negative (Pα(-)S) populations were sorted by flow cytometry. B) PDGFRA-positive (Pα(+)S) and PDGFRA-negative (Pα(-)S) MSCs were latently infected with rKSHV.219 and analyzed for GFP expression using a fluorescence microscope. C) Immunofluorescence analysis of KSHV-infected Pα(+)S (K-Pα(+)S) and Pα(-)S (K-Pα(-)S) to evaluate KSHV LANA expression (red), nuclei were counterstained with DAPI (blue). D) Fold-changes in KSHV gene expression between 24 hours KSHV post-infection and after KSHV latency establishment in MSC or KS-like media as determined by RT-qPCR. Triplicates are shown as means ± SD. K-Pα(-)S population on the top and K-Pα(+)S population on the bottom. *P < 0.05. E) Soft agar colony formation assay to determine anchorage-independent cell growth in Pα(+)S KS, K-Pα(+)S MSC, and K-Pα(+)S KS cells. F) Tumor-Free mice curve from subcutaneous injection of Pα(+)S KS, K-Pα(+)S MSC, and K-Pα(+)S KS cells into nude mice. N=6. G) Representative microscopic histological section of mouse KS-like mECK36 tumor and K-Pα(+)S KS tumor stained with hematoxylin and eosin (H&E). H) Immunofluorescence analysis of KSHV LANA (red) in K-Pα(+)S KS tumor, nuclei were counterstained with DAPI (blue). I) Immunofluorescence analysis of PECAM1 (red) in K-Pα(+)S KS tumor, nuclei were counterstained with DAPI (blue).

The cellular mechanisms promoting inflammation, wound repair and angiogenesis, may promote the development of KS tumors in KSHV-infected individuals. To test the effect of a pro-angiogenic KS-like environment on KSHV tumorigenesis, we infected mouse bone marrow-derived PDGFRA-positive (Pα(+)S) and PDGFRA-negative (Pα(-)S) MSCs with rKSHV.219. We then incubated the cells in KS-like media (K-Pα(+)S KS), a media that is rich in heparin and endothelial cell growth factors (ECGF), which reproduces a pro-angiogenic/vasculogenic environment in the culture [41, 42]. The percentage of KSHV *de novo* infection was similar (80%) in both MSC and KS condition as well as in PDGFRA-negative and PDGFRA-positive cells (S2 Figure). After two weeks of puromycin selection, we obtained KSHV-infected PDGFRA-negative (K-Pα(-)S KS) or PDGFRA-positive MSCs growing in KS-like media (K-Pα(+)S KS). We compared KSHV gene expression in K-Pα(-)S and K-Pα(+)S growing in KS-like media with those maintained and selected in MSC media. Interestingly, we found upregulation of KSHV latent and lytic genes (LANA, RTA, vGPCR, and vIRF1) only in K-Pα(+)S cells propagated in KS-like culture conditions (Figure 1D), indicating that this cell lineage and environmental conditions favor KSHV gene expression *in vitro*.

### KSHV infection is tumorigenic only in PDGFRA-positive MSCs (Pα(+)S) grown in pro­angiogenic KS-like environmental conditions

Since we showed that pro-angiogenic KS-like culture conditions favor KSHV gene expression *in vitro* (Figure 1D) including many viral lytic oncogenes, we determined whether these conditions also conferred malignant cell characteristics to KSHV-infected Pα(+)S MSCs, such as anchorage-independent cell growth, we performed a soft-agar colony 9 formation assay. Uninfected PDGFRA-positive MSCs cultured in KS-like media and KSHV-infected PDGFRA-positive MSCs cultured in MSC media did not grow on soft-agar (Figure 1E). In contrast, KSHV-infected PDGFRA-positive (Pα(+)S) MSCs cultured in pro-angiogenic KS-like conditions (K-Pα(+)S KS) were the only cells to form colonies in soft agar (Figure 1E), suggesting that the KS-like culture conditions conferred malignant phenotypic characteristics to K-Pα(+)S MSCs. More importantly, subcutaneous injection of K-Pα(+)S KS cells into nude mice resulted in tumor formation in all 6 injected mice by 7 weeks. No tumors formed from either uninfected Pα(+)S MSC growing in KS-like culture conditions or in infected Pα(+)S growing in MSC culture conditions (Figure 1F). Paraffin-embedded sections of KSHV-infected PDGFRA-positive MSCs tumors (K-Pα(+)S KS tumor) stained with hematoxylin and eosin (H&E) were analyzed by a pathologist in a blind manner, confirming that these tumors were indistinguishable from the vascularized spindle cell sarcomas formed by mECK36 tumors previously generated by our lab, which were thoroughly characterized as KS-like tumors [26] (Figure 1G). Using immunofluorescence detection of the KSHV LANA protein, we confirmed that the majority of the tumor cells display the classic punctate nuclear staining (Figure 1H). Additionally, using the endothelial cell marker PECAM1, we observed that these tumors displayed phenotypic markers that corresponded to those typically found in human KS lesions (Figure 1I). We concluded that KSHV infection of PDGFRA-positive MSCs in combination with a pro-angiogenic KS-like environmental condition induces a malignant transformed phenotype in these cells, leading to tumorigenicity in nude mice.

### Upregulation of KSHV lytic gene expression during *in vivo* tumorigenesis

*We* have previously shown that KSHV tumorigenesis in the mECK36 mouse KS-like model occurs with concomitant up-regulation of KSHV lytic genes and angiogenic ligands/receptors [14, 26]. This KSHV *in vivo* lytic switch is thus responsible for the activation of many autocrine and paracrine oncogenic signaling cascades, including the recently reported ligand-mediated activation of PDGFRA signaling pathway that drives KS tumorigenesis [14]. RNA-sequencing (RNA-seq) analysis of KSHV gene expression in tumors derived from KSHV-infected PDGFRA-positive MSCs grown in KS-like media (K-Pα(+)S KS tumors), which were found to be histologically indistinguishable from mECK36, also showed upregulation of KSHV lytic gene expression compared to the tumorigenic cells grown *in vitro* (KSHV *in vivo* lytic switch) (Figure 2A). We also compared the KSHV transcriptomes of our infected cells and tumors with those obtained from published RNA-seq analysis of actual Kaposi’s Sarcoma human biopsies [43] from AIDS-KS patients (Figure 2B-C). As shown in Figure 2B, the unsupervised hierarchical clustering analysis show that human KS samples cluster between the lytic-expressing mouse KS-like tumors and the latently infected K-Pα(+)S KS cells. This shows that the KSHV transcriptomes of MSCs grown in KS-like conditions are closer to those of actual human KS tumors than those grown in MSC conditions. They also show that the transcriptomes of the MSCs tumors are representative of some human KS samples as we previously reported for the mECK36 model [26]. In fact, some human KS samples showed upregulation of KSHV lytic genes when compared with the latently infected mouse MSCs grown in either MSC or KS-like conditions, further supporting the relevance of KSHV *in vivo* lytic gene expression observed in MSCs tumors (the *in vivo* lytic switch) to actual cases of human AIDS-KS. The upregulation of KSHV lytic genes in K-Pα(+)S KS tumors correlated with PDFGRA signaling activation (Figure 2D) and phospho-PDGFRA co-distributed with KSHV LANA (Figure 2E), further supporting our investigated link between KSHV and PDGFRA activation in these tumors [14]. Host gene RNA-seq comparison of KSHV-infected PDGFRA-positive cells grown in KS-like media (K-Pα(+)S KS) to their corresponding tumors in nude mice revealed 1,861 differentially expressed genes (DEGs) (Figure 2F). The analysis of these DEGs using Gene Ontology (GO), Kyoto Encyclopedia of Genes and Genomes (KEGG) and Reactome databases, uncovered major changes in pathways involving glycolysis, respiratory electron transport, NOTCH signaling, TGFB signaling and antigen processing and presentation among the most up-regulated pathways during KSHV *in vivo* lytic switch and tumorigenesis (Figure 2G). The results from these bioinformatics assessments are consistent with prior biological studies on several KSHV lytic genes and lytic conversions upon environmental conditions [13].

**Figure 2.**
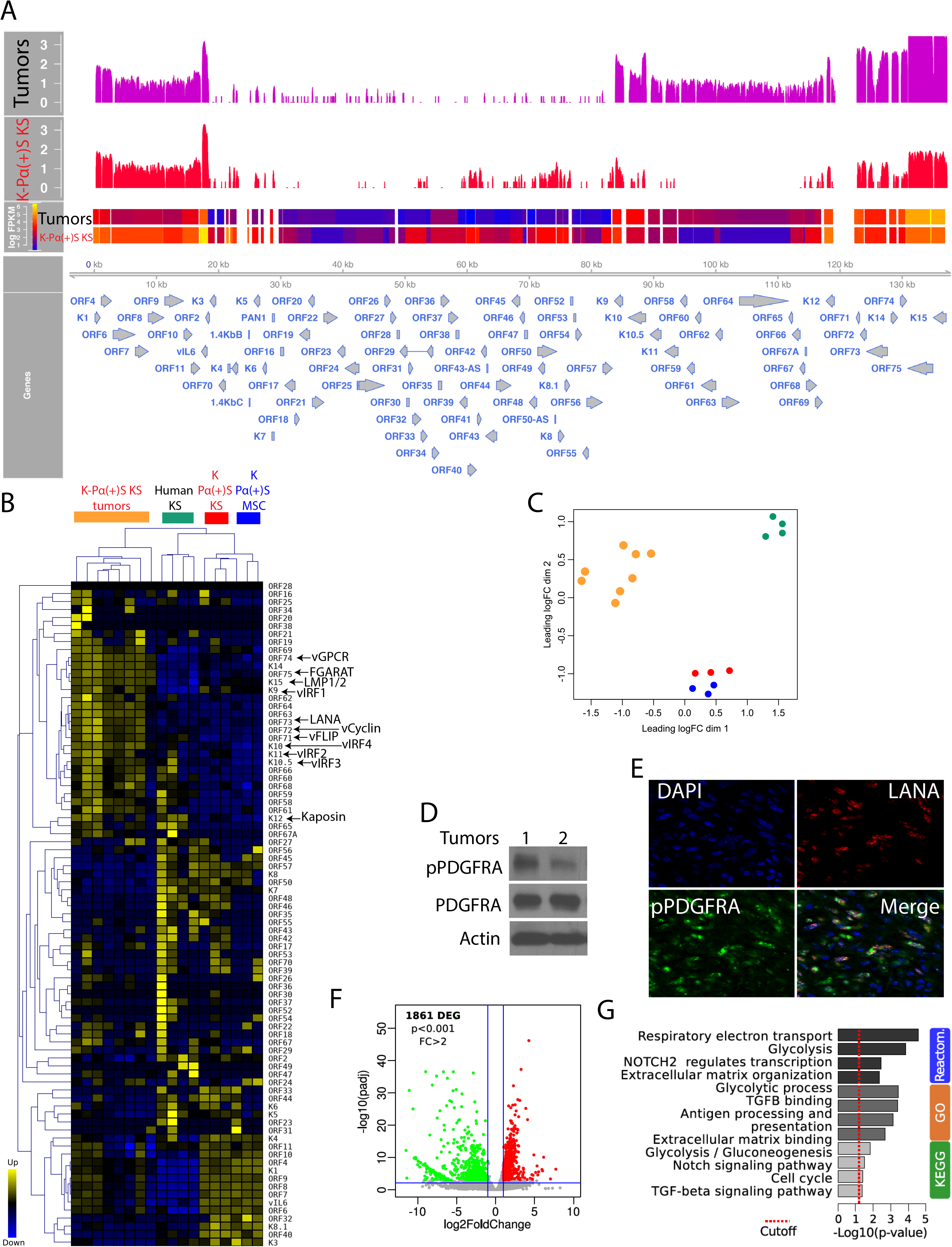
Upregulation of KSHV lytic gene expression during *in vivo* tumorigenesis. A) Genome-wide analysis of KSHV transcripts by RNA deep sequencing, comparison of the transcription profiles of K-Pα(+)S KS cells (N=3) and K-Pα(+)S KS tumors (N=8) derived from these cells. Transcriptional levels of viral genes were quantified in reads per kilobase of coding region per million total read numbers (RPKM) in the sample. The y-axis represents the number of reads aligned to each nucleotide position and x-axis represents the KSHV genome position. B) Unsupervised hierarchical clustering of KSHV transcriptome among K-Pα(+)S KS cells, K-Pα(+)S KS tumors and human KS tumors. The arrows indicate KSHV oncogenes. C) Multidimensional scaling plot showing the distance of each sample from each other determined by their KSHV expression profiles. D) Total and phospho-PDGFRA were determined by immunoblotting in two MSC K-Pα(+)S KS tumors. Actin was used as a loading control. E) Immunofluorescence analysis of K-Pα(+)S KS tumors to detect KSHV LANA (red) and phospho-PDGFRA (green). Cell nuclei were counterstained with DAPI (blue). F) Volcano plot showing 1,861 differentially expressed genes (DEGs) analyzed by RNA-Seq between K-Pα(+)S KS tumors *in vivo* and K-Pα(+)S KS cells *in vitro*. G) Functional enrichment analysis based on genes differentially expressed among K-Pα(+)S KS tumors and K-Pα(+)S KS cells.

These data also agree with the predictions from the analysis of a human KS-signature and newer RNA-sequencing studies derived from differentially expressed genes between KS and normal skin, which predicted the activation of several paracrine axes by upregulation of receptors and/or their ligands [2] and changes in TGFB signaling and glucose metabolism [43] similar to those observed in our MSC tumors.

### A de-repressed KSHV epigenome allows for expression of oncogenic KSHV genes in tumorigenic PDGFRA-positive MSCs growing in a KS-like environment

To understand the differences in oncogenicity between KSHV-infected PDGFRA-positive MSCs grown in MSC versus KS-like media (K-Pα(+)S KS versus K-Pα(+)S MSC); we analyzed the pattern of KSHV viral gene expression using RNA-seq analysis (Figure 3A). The results from this analysis were in line with the qRT-PCR results depicted in figure 1D which showed increased KSHV gene expression in cells grown in KS-like media (K-Pα(+)S KS), and also revealed an overall increased expression of KSHV genes in these tumorigenic cells.

**Figure 3.**
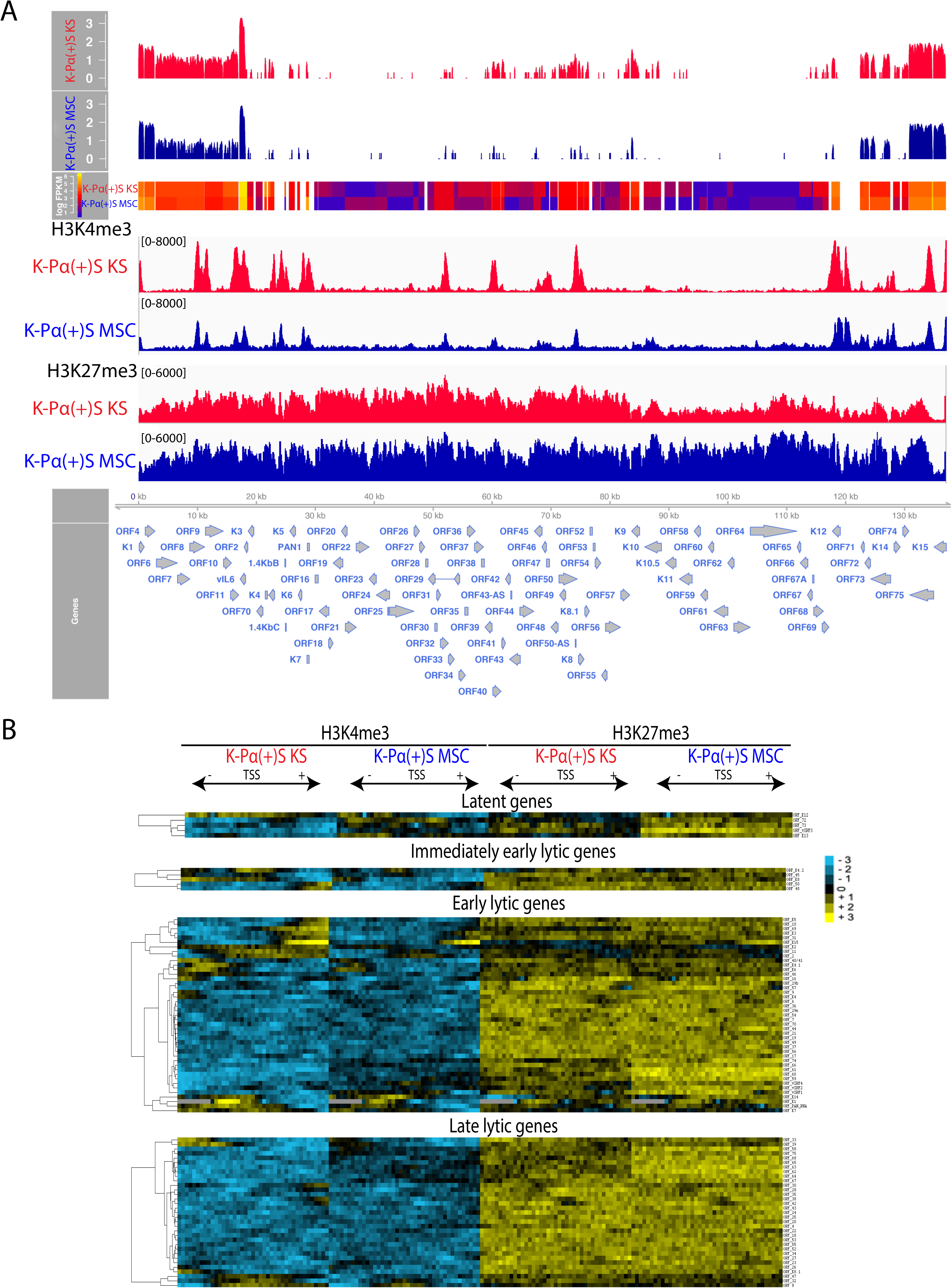
**A de-repressed KSHV epigenome allows for expression of oncogenic KSHV genes in tumorigenic PDGFRA-positive MSCs growing in KS-like environment.** A) Genome-wide analysis of KSHV transcripts by RNA deep sequencing, comparison of the transcription profiles of MSC K-Pα(+)S MSC and K-Pα(+)S KS cells. Transcriptional levels of viral genes were quantified in reads per kilobase of coding region per million total read numbers (RPKM) in the sample. The y-axis represents the number of reads aligned to each nucleotide position and x-axis represents the KSHV genome position. Global patterns of H3K4 tri-methylation (HgK4me3) and H3K27 tri-methylation (H3K27me3) in K-Pα(+)S MSC and K-Pα(+)S KS cells were analyzed by ChIP-seq assays as described in the text. Values shown on the y-axis represent the relative enrichment of normalized signals from the immunoprecipitated material over input. B) Heat map representation of changes in histone modifications at the gene regulatory regions of KSHV genes grouped by expression class Latent, Immediate-early lytic, Early lytic and Late lytic genes. The rows display the relative abundance of the indicated histone modification within the −1 kb to +1 kb genomic regions flanking the translational start site (TSS) of each viral gene. The blue and yellow colors denote lower-than-average and higher-than-average enrichment, respectively.

Importantly, we verified that these differences in viral gene expression are not due to differences in viral DNA copy number (S3 Figure). Many of these KSHV genes are well-characterized viral oncogenes such as vGPCR (ORF74), K12, vIRF1 (ORFK9), vFLIP (ORF71), K15. Our data suggest that PDGFRA-positive KSHV MSCs growing in a pro-angiogenic KS-like environment (K-Pα(+)S KS) display a unique KSHV transcriptional profile that encompasses enhanced expression of oncogenic KSHV genes, without KSHV completing its full viral lytic cycle (abortive lytic state), which likely leads to KSHV-induced tumorigenicity. Several studies indicate that histone modifications play a role in the epigenetic regulation of KSHV gene expression [44, 45]. Activating histone modifications like H3K4me3 are enriched in some activated loci while repressive histone modifications like H3K27me3 are widespread across the KSHV viral genome [46–48]. We determined if the differences in KSHV viral gene expression between non-tumorigenic K-Pα(+)S MSC and tumorigenic K-Pα(+)S KS cells could be attributed to differential epigenetic regulation at the viral promoters. We performed chromatin immunoprecipitation followed by next-generation sequencing (ChlP-seq) in three biological replicates to map the distribution of two histone modifications, H3K4me3, and H3K27me3, that are enriched at actively transcribed and repressed genes, respectively [49]. We found that KSHV chromatin contained both H3K4me3 and H3K27me3 marks; however, with distinct patterns of association at KSHV genomic regions between K-Pα(+)S KS versus K-Pα(+)S MSC cells (Figure 3A). As previously reported, the latency-associated locus where KSHV genes that are constitutively expressed during latency are located, was enriched with H3K4me3 in both K-Pα(+)S KS and K-Pα(+)S MSC cells (Figure 3A) [46, 48]. Interestingly, H3K27me3 was widely distributed throughout the KSHV genome in both cells populations, but this repressive mark was more enriched in the KSHV genome in non-tumorigenic K-Pα(+)S MSC cells. This result correlates with the less KSHV gene expression observed in these cells by qRT-PCR and RNA-seq. On the other hand and in agreement with the KSHV gene expression upregulation seen in tumorigenic K-Pα(+)S KS cells, we observed more H3K4me3 activating marks on the KSHV genome of these tumorigenic cells, especially in immediate early and early lytic genes (Figure 3A).

To delineate the characteristics of the chromatin structures associated with the regulatory regions of KSHV genes, we aligned the KSHV open reading frames (ORFs) relative to their translational start sites (TSS). We plotted the signal intensities of probes derived from the ChIP-Seq analysis across a 2 kb region spanning 1 kb on either side of the TSS (please see Materials and Methods for details) [46] (Figure 3B). Since RTA is responsible for the switch between latency and lytic replication, its promoter is not only tightly repressed during latency, but its silencing should also be rapidly reversible upon reactivation. We found that the RTA promoter is enriched in both H3K4me3 and H3K27me3 during latency (Figure 3 A-B), suggesting it possesses a bivalent chromatin that maintains repression of the RTA promoter while keeping it poised for rapid activation, in agreement with what was previously reported using BCLBC-1, TRExBCBL1-RTA and SLK cells [46, 47]. Moreover, in K-Pα(+)S KS cells the RTA promoter and other immediately early and early lytic promoters showed more activating marks than in K-Pα(+)S MSC cells. These results point to epigenetic regulation as a contributory mechanism for the unique KSHV transcriptional program shown by K-Pα(+)S KS tumorigenic cells *in vitro* and support our MSC mouse model for studying the connection between epigenetic regulation of KSHV gene expression and oncogenesis.

### Global gene expression profiling and histone mark distribution in K-Pα(+)S KS and K-Pα(+)S MSC cells

We showed that KSHV gene expression differences between non-tumorigenic K-Pα(+)S MSC and tumorigenic K-Pα(+)S KS cells are explained at least, in part, by differences in epigenetic regulation of the KSHV genome. In order to compare KSHV with host gene expression regulation in these two cell lines, we carried out a global gene expression profile by RNA-seq analysis. We found more than 450 differentially expressed host genes (DEGs) between tumorigenic K-Pα(+)S KS and non-tumorigenic K-Pα(+)S MSC cells (Figure 4A). These DEGs were enriched for multiple biological processes; notably, PDGFR signaling, PI3K-AKT and P53 signaling, together with cell cycle regulation (Figure 4B). Moreover, and as we showed for KSHV gene expression (Figures 3), H3K27me3 and H3K4me3 ChIP-seq revealed slightly more H3K4me3 enrichment (Figure 4C) and fewer H3K27me3 (Figure 4D) at the promoters of these upregulated genes in tumorigenic K-Pα(+)S KS cells, further confirming epigenetic regulation of transcription at these loci.

**Figure 4.**
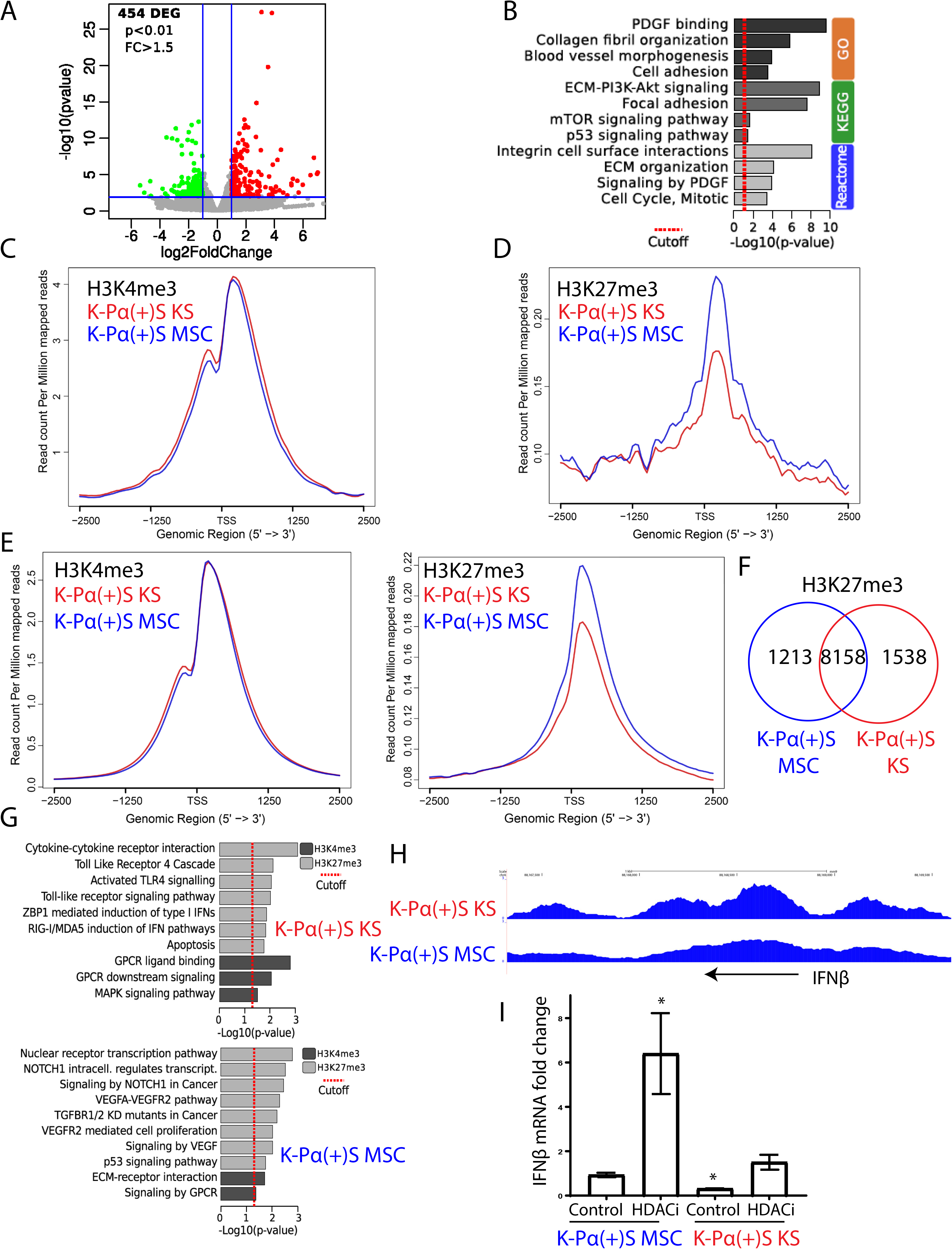
Global gene expression profiling and histone marks distribution in K-Pα(+)S KS and K-Pα(+)S MSCs. A) Volcano plot showing 454 differentially expressed genes (DEGs) analyzed by RNA-seq between K-Pα(+)S MSC and K-Pα(+)S KS cells. B) Functional enrichment analysis based on genes differentially expressed among K-Pα(+)S MSC and K-Pα(+)S KS cells. C) H3K4me3 ChIP-seq signals on DEGs in K-Pα(+)S MSC and K-Pα(+)S KS cells. D) H3K27me3 ChIP-seq signals on DEGs in K-Pα(+)S MSC and K-Pα(+)S KS cells. E) H3K4me3 (left) and H3K27me3 (right) ChIP-seq signals in K-Pα(+)S MSC and K-Pα(+)S KS cells. F) Venn diagrams showing the overlap between H3K27me3-enriched genes in K-Pα(+)S MSC and K-Pα(+)S KS cells. G) Pathway analysis representing KEGG and REACTOME pathway enriched by the genes with differentially enriched regions (DERs) of H3K27me3 and H3K4me3 in K-Pα(+)S MSC (bottom) and K-Pα(+)S KS (top) cells. H) Genome browser screenshots of H3K27me3 ChIP-seq signal in K-Pα(+)S MSC and K-Pα(+)S KS cells at the IFNP genomic region. I) Fold-changes of IFNP gene expression in K-Pα(+)S MSC and K-Pα(+)S KS cells before and after KSHV lytic reactivation, determined by RT-qPCR. Triplicates are shown as means ± SD. *P < 0.05.

Interestingly, Chip-seq analysis for the distribution of the two histone modifications, H3K4me3 and H3K27me3, over the host genome revealed that K-Pα(+)S MSC cells displayed more global enrichment of H3K27me3 at host genes compared with K-Pα(+)S KS cells, suggesting that the host genome of K-Pα(+)S MSC cells is more transcriptionally repressed genome-wide (Figure 4E). To characterize these repressed genes, we associated H3K27me3 ChIP-seq peaks to all the genes of the mouse genome. We found that while the number of genes decorated with H3K27me3 in the two K-Pα(+)S cells were comparable (9,696 in KSHV KS cells versus 9,371 in K-Pα(+)S MSC cells) with a large number of genes common between the two (8,158 genes decorated with H3K27me3 in both), 1,538 genes were exclusive to K-Pα(+)S KS cells while 1,213 were exclusive to K-Pα(+)S MSC cells (Figure 4F). KEGG and REACTOME pathway enrichment analysis was performed for these repressed H3K27me3 gene targets that are specific to K-Pα(+)S KS or K-Pα(+)S MSC. In K-Pα(+)S MSC cells the most repressed genes involved NOTCH1 regulation, signaling by NOTCH1 in Cancer, VEGFA-VEGFR2 pathway, TGFBR1/2 KD mutants in cancer, VEGFR2-mediated cell proliferation, signaling by VEGF and P53 signaling pathways (Figure 4G). These aforementioned genes all related to KS and KSHV oncogenic signaling, which thus, appear more de-repressed in the oncogenic K-Pα(+)S KS cells.

Interestingly, in the K-Pα(+)S KS cells, the most repressed genes involved Toll-Like Receptor 4 cascade, Activated TLR4 signaling, Toll-like receptor signaling pathway, ZBP1-mediated induction of type 1 IFNs and RIG-1/MDA5 induction of IFN pathways, all related to the innate immune response (Figure 4G-H) and the Type I IFN (IFNα and β) anti-viral response. This suggests that the IFN anti-viral response would be more repressed in KSHV-infected cells grown in a KS-like environment. In fact, K-Pα(+)S MSC cells showed higher expression of IFNβ mRNA than K-Pα(+)S KS cells confirming the repression of the *IFN*β loci in K-Pα(+)S KS cells (Figure 4I). Moreover, we found that only K-Pα(+)S MSC cells upregulated *IFN*β expression after KSHV lytic reactivation with an HDAC inhibitor (SAHA/ Vorinostat) (Figure 4I). Taken together our RNA-seq and ChIP-seq analysis on host and KSHV genes indicates that tumorigenic K-Pα(+)S KS cells are more adapted to withstand oncogenic KSHV lytic gene expression possibly by regulating PDGFR, PI3K-AKT, P53 and cell cycle pathways, but also by repressing innate immune response genes upon induction of the lytic replication program.

### A permissive epigenetic landscape induced by a pro-angiogenic environment promotes an oncogenic viral lytic-driven mechanism of tumorigenesis

Our results suggest that KSHV and host epigenetic mechanisms which regulate KSHV lytic gene expression dictate at least, in part, the differences in tumorigenicity between cells grown in MSC and KS media. Figure 2A shows that K-Pα(+)S KS tumorigenicity occurs along with a steep increase in KSHV lytic gene expression (KSHV *in vivo* lytic switch). In order to model the biological consequences of this tumorigenic lytic switch in K-Pα(+)S KS and K-Pα(+)S MSC cells in an *in vitro* system, we used HDAC inhibitors (HDACi) such as SAHA/ Vorinostat which we and others have shown are powerful inducers of KSHV lytic replication in several systems [50–53].

We found that K-Pα(+)S MSC cells, which are not tumorigenic, massively upregulated KSHV lytic genes (100-folds) after 24 hours of SAHA treatment, as shown using qRT-PCR (Figure 5A). In contrast, in tumorigenic K-Pα(+)S KS cells that display a higher level of basal KSHV lytic gene expression, the fold-increase expression of KSHV lytic genes after reactivation was an order of magnitude lower than in K-Pα(+)S MSC cells (Figure 5A). We found that KSHV reactivation following SAHA treatment induces the appearance of a RFP (early lytic marker) positive cell population in both K-Pα(+)S MSC (up to 10%) and K-Pα(+)S KS (up to 3%) cells (Figure 5B). To characterize the pattern of KSHV gene expression in GFP-positive and RFP-positive cell populations generated after reactivation, we used fluorescence-activated cell sorting (FACS) to separate these two populations. We found that the RFP positive populations of both K-Pα(+)S MSC and K-Pα(+)S KS cells (Figure 5C, left panel) showed massive upregulation of KSHV lytic and latent genes, when compared to un­induced cells, characteristic of lytic induction. The GFP-positive/RFP-negative population from the non-tumorigenic K-Pα(+)S MSC cells also showed upregulation of both KSHV latent and lytic genes (LANA, vFLIP and vCyclin, RTA, vGCPR, K8.1) after SAHA treatment, suggesting KSHV lytic reactivation (Figure 5C, right panel) in this population as well. On the other hand, the GFP-positive/RFP-negative population from the tumorigenic K-Pα(+)S KS cells showed only upregulation of some specific lytic genes (RTA and K8.1) (Figure 5C, right panel). Thus, the RFP-lytic marker distribution allowed us to identify a population of “classic” lytically induced cells that are similar in both K-Pα(+)S MSC and K-Pα(+)S KS cells. Interestingly, the GFP-positive/RFP-negative population, which corresponds to the vast majority of the cells, shows a distinct pattern of KSHV gene expression between K-Pα(+)S MSC and K-Pα(+)S KS cells after KSHV lytic reactivation. These results suggest that differences in the epigenetic environment induced by KS-like culture conditions can lead to a different outcome for lytic reactivation in K-Pα(+)S KS cells and more importantly, be a key determinant for the oncogenic consequences of the KSHV lytic switch.

**Figure 5.**
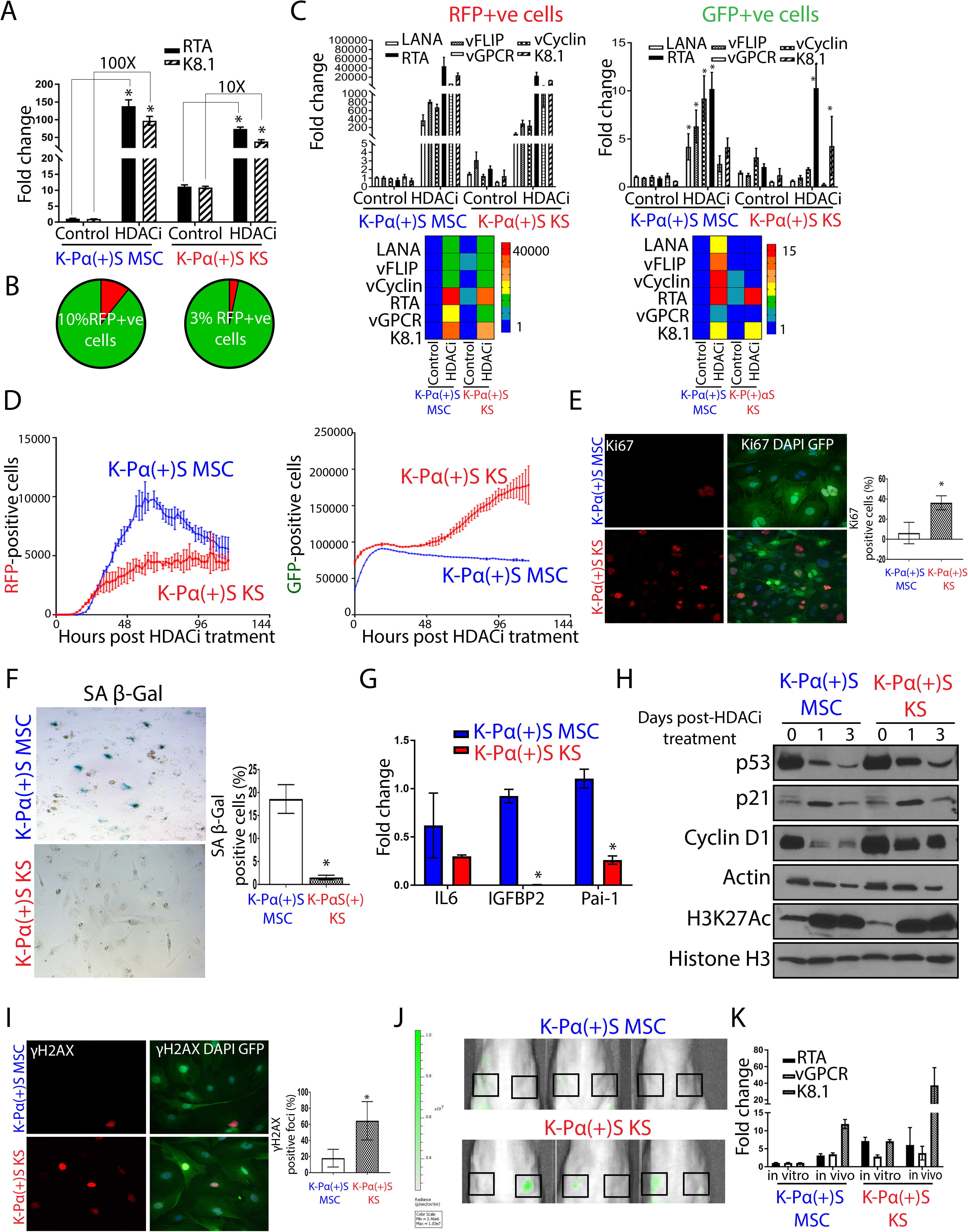
**A permissive epigenetic landscape induced by a pro-angiogenic environment promotes an oncogenic viral lytic-driven mechanism of tumorigenesis.** A) Fold-changes of KSHV gene expression in K-Pα(+)S MSC and K-Pα(+)S KS cells after SAHA (7/5 uM) treatment for 24 hours, determined by RT-qPCR. Triplicates fold change to un­induced K-Pα(+)S MSC are presented as means ± SD. *P < 0.05. B) Percentage of RFP-positive and GFP-positive cells after 72 hours of KSHV reactivation measured by Flow cytometry in K-Pα(+)S MSC and K-Pα(+)S KS cells. C) Left panel, fold-changes of KSHV gene expression in sorted RFP-positive population of K-Pα(+)S MSC and K-Pα(+)S KS cells after 72 hours of SAHA treatment were determined by RT-qPCR. Triplicates fold change to un-induced K-Pα(+)S MSC are presented as means ± SD. *P < 0.05. The heatmap for the mean RT-qPCR data is shown below. Right panel, fold-changes of KSHV gene expression in sorted GFP-positive population in K-Pα(+)S MSC and K-Pα(+)S KS cells after 72 hours of SAHA treatment were determined by RT-qPCR. Triplicates fold change to un-induced K-Pα(+)S MSC are presented as means ± SD. *P < 0.05. The heatmap for the mean RT-qPCR data is shown below. D) K-Pα(+)S MSC and K-Pα(+)S KS cells were treated with SAHA and incubated in an Incucyte Zoom (Essen Bioscience), acquiring red and green fluorescence images. The cell proliferation graphs of RFP-positive (left panel) and GFP-positive (right panel) cells were graphed overtime as a measure of proliferation and shown as mean ± SE from three replicates for each condition. E) Immunofluorescence analysis of K167 expression (red) was performed after 72 hours of SAHA treatment on GFP-positive K-Pα(+)S MSC and GFP-positive K-Pα(+)S KS cells; nuclei were counterstained with DAPI (blue).The quantification is shown at the right of the images. Values are presented as means ± SD. *P < 0.05. F) SA *β*-gal staining was performed after 72 hours of SAHA treatment on K-Pα(+)S MSC and K-Pα(+)S KS cells. The quantification is shown at the right of corresponding panels. Values are presented as means ± SD. *P < 0.05 G) mRNA fold-changes of markers of senescence and SASP or Senescence Associated Secretory Phenotype in K-Pα(+)S MSC and K-Pα(+)S KS after 72 hours of SAHA treatment were assessed by RT-qPCR in triplicate and are presented as means ± SD. *P <0.05. Η) ρ53, Ρ2ΐ, and Cyclin Di levels were analyzed by immunoblotting in K-Pα(+)S MSC and K-Pα(+)S KS cells after 72 hours of SAHA treatment. Histone 3 K27 Acetylation (H3K27AC) was used as the control for HDAC inhibition by SAHA. Actin was used as loading control. I) Immunofluorescence analysis of I2IH2AX expression (red) was performed after 72 hours of SAHA treatment on GFP-positive K-Pα(+)S MSC and GFP-positive K-Pα(+)S KS cells; nuclei were counterstained with DAPI (blue). The quantification is shown at the right of the images. Values are presented as means ± SD. *P < 0.05. J) Spectrum In Vivo Imaging System (IVIS) of Matrigel-plugs (1 week after injection) containing GFP-positive PDGFRA-positive MSCs from K-Pα(+)S MSC and K-Pα(+)S KS cells. Black Squares indicate site of injection. K) Fold-changes of KSHV gene expression in K-Pα(+)S MSC and K-Pα(+)S KS after 1 week *in vivo* Matrigel-plug were assessed by RT-qPCR in triplicate and are presented as means ± SD.

To determine if differences in the culture conditions can affect the proliferation of infected cells upon the lytic switch, we used an IncuCyte® Live-Cell Imaging and Analysis System, which enables real-time, automated cell proliferation assays to follow the growth of RFP-positive and GFP-positive populations among SAHA-induced cells. K-Pα(+)S MSC and K-Pα(+)S KS RFP-positive populations, which we found, do not differ in their pattern of KSHV induction (Figure 5D, left panel), augment in cell number upon lytic induction, then reach a limit of cell growth. In contrast, in the case of GFP-positive cells, we found that the K-Pα(+)S MSC cells stop proliferating after reactivation (Figure 5D, right panel), while induced K-Pα(+)S KS cells continue to proliferate. Interestingly, Figure 5C shows that both cells display KSHV lytic gene expression, but only the tumorigenic K-Pα(+)S KS cells are able to continue proliferating in spite of lytic KSHV gene expression. Moreover, K-Pα(+)S KS cells showed more proliferation markers (such as Ki67) than K-Pα(+)S MSC cells (Figure 5E) after 72hS of KSHV lytic reactivation.

Cell cycle arrest and cellular senescence are known to occur during the lytic replication phase of herpesviruses and other DNA viruses [54]. This type of senescence, characterized by the upregulation of cell cycle inhibitors such as P53 and p21, is generally also induced by aberrant proliferative signals of oncogenes such as activated Ras (RasV12) leading to increased oxidative DNA damage [55, 56]. The increase in expression of many oncogenic KSHV genes occurring during lytic replication could induce oncogenic stress that would typically be expected to trigger oncogene-induced senescence (OIS) [57, 58]. A widely used assay for cell senescence is the cytochemical detection of acidic β-galactosidase activity (pH 6.0), termed senescence-associated β-galactosidase (SA-β-Gal) [59]. In order to fulfill all the hallmarks for establishing cell senescence [60, 61] we measured 1) Markers for Inhibition of the cell cycle and levels of cell cycle arrest proteins 2) Expression of acidic beta-galactosidase and 3) Upregulation of Senescence Associated Secretory Phenotype (SASP) and other senescence markers in K-Pα(+)S MSC and K-Pα(+)S KS cells after KSHV reactivation. As shown in Figure 5E-H, K-Pα(+)S MSC cells: 1) Displayed decreased levels of proliferation markers such as Ki67 and expression of cyclin D1 2) Displayed enhanced features and markers of cell senescence, such as increased SA-β-Gal staining, and upregulation of SASP and senescence markers (IL-6, IGFBP2, PAI-1), 3) Expressed cell cycle inhibitors such P53 and p21 that correlated with their arrest of cellular proliferation and increase in the number of cells that underwent senescence. On the other hand, tumorigenic K-Pα(+)S KS cells showed low SA-β-Gal staining, lower levels of SASP and senescence markers, higher expression of Ki67 and upregulated cyclin D1, despite upregulation of p21 and P53 after lytic reactivation, which correlated with their increase in proliferation (Figure 5D-H). Importantly, K-Pα(+)S MSC cells showed less DNA damage response (such as H2AX) (Figure 5I), compared to tumorigenic K-Pα(+)S KS cells after KSHV reactivation. These tumorigenic cells display robust proliferation markers in spite of an increased level of DNA repair foci, further indicating that these cells are able to overcome KSHV-driven oncogene-induced senescence as well as DNA damage-induced cell cycle arrest, continuing to proliferate after KSHV reactivation.

In order to study the consequences of the *in vivo* KSHV lytic switch in more physiological conditions, we performed a short-term *in vivo* Matrigel-plug experiment with K-Pα(+)S MSC and K-Pα(+)S KS cells. Within the Matrigel-plug, the cells were subjected to *in vivo* growth conditions fora period of one week. We injected K-Pα(+)S MSC and K-Pα(+)S KS cells subcutaneously into Matrigel in nude mice, and after one week of *in vivo* growth, we analyzed the growth of viable cells by the GFP-expression using Spectrum In Vivo Imaging System (IVIS). We also extracted the Matrigel-plug containing cells to document the *in vivo* KSHV gene expression lytic switch by qRT-PCR. After one week of *in vivo* growth, K-Pα(+)S KS cells continued proliferating, as shown by the GFP-positive signal at the site of injection, however, K-Pα(+)S MSC cells were not able to proliferate *in vivo,* as seen by the lack of GFP signal (Figure 5-1). Real time qRT-PCR analysis showed that both cell lines displayed upregulation of KSHV lytic genes (Figure 5K); yet, only tumorigenic K-Pα(+)S KS cells were able to proliferate in the context of the KSHV *in vivo* lytic switch. On the other hand, the growth inhibition shown by K-Pα(+)S MSC cells directly occurred together with the upregulation of KSHV gene expression *in vivo* (Figure 5K). The results obtained with this Matrigel-plug in *in vivo* experiments mimic those observed with KSHV lytic induction *in vitro* (Figure 5D). Taken together, these *in vitro* and *in vivo* results demonstrate that PDGFRA-positive KSHV-infected MSCs are able to proliferate *in vitro* and *in vivo* in the presence of KSHV oncogenic gene expression only when they are subjected to a pro-angiogenic KS-like environment. Thus, these conditions not only conferred an epigenetic adaptation leading to increased KSHV gene expression but also should confer a mechanism, which upon *in vivo* lytic switch, will allow them to overcome KSHV-driven oncogene-induced senescence and cell cycle arrest, leading to tumorigenesis.

### PDGFRA signaling allows KSHV-infected PDGFRA-positive MSCs grown in a KS-like environment to continue proliferating after lytic reactivation

We sought to elucidate the mechanism that allows K-Pα(+)S KS cells to continue proliferating after KSHV lytic reactivation. To test, in an unbiased manner, which tyrosine kinase receptors were differentially activated between K-Pα(+)S MSC and K-Pα(+)S KS cells after lytic reactivation, we used a tyrosine kinase proteomic array (Figure 6A). We found that PDGFRA was the most differentially activated tyrosine kinase over 39 RTKs tested in the array (Figure 6A). This finding was consistent with the pathway enrichment analysis shown in Figure 4B, K-Pα(+)S cells grown in KS-like media display enhanced expression of a genetic network enriched in genes that regulate PDGFR signaling, AKT and cell cycle. To characterize the involvement of PDGFRA activation and downstream signaling pathways, we analyzed the cells after KSHV reactivation for PDGFRA and AKT signaling activation. K-Pα(+)S KS cells showed activation of both PDGFRA and AKT signaling compared to K-Pα(+)S MSC cells (Figure 6B). Moreover, we found that activation of PDGFRA signaling in induced K-Pα(+)S KS cells correlated with upregulation of its ligands PDGFA and PDGFB (Figure 6C), further confirming the fact that KSHV lytic expression induces ligand-mediated activation of this oncogenic driver [14]. This KSHV-induced ligand-mediated activation of PDGFRA signaling could constitute an important mechanism mediating the adaptation of the K-Pα(+)S KS cells to continue proliferating after lytic reactivation.

**Figure 6.**
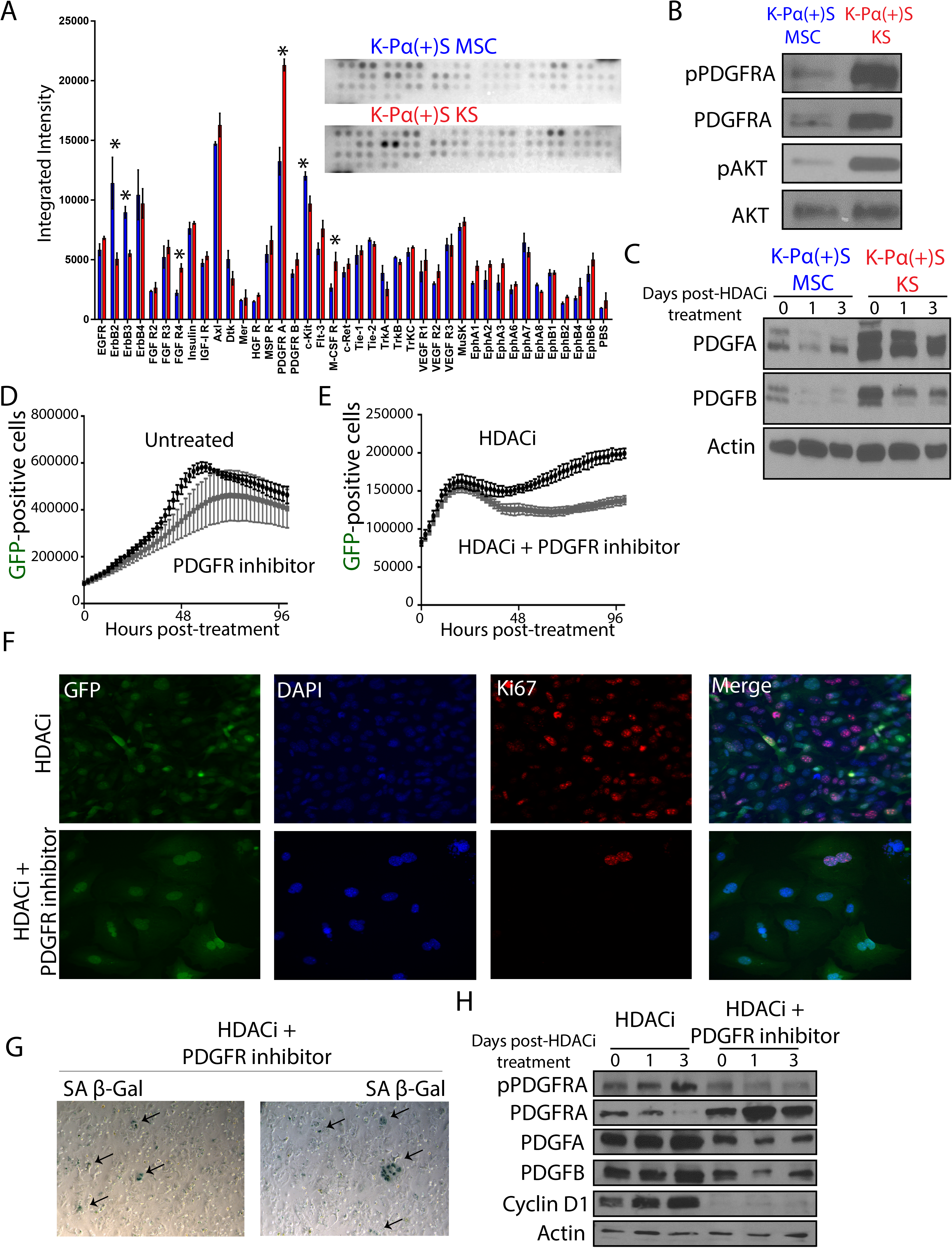
**PDGFRA signaling allows KSHV-infected PDGFRA-positive MSCs grown in KS-like environment to continue proliferating after lytic reactivation.** A) Mouse Phospho-Receptor Tyrosine Kinase (RTK) Array Kit used to quantify levels of phosphorylation of 39 RTKs in K-Pα(+)S MSC and K-Pα(+)S KS cells after 72 hours of SAHA treatment pointing to major activation spot corresponding to PDGF receptor alpha chain. B) Total and phospho-PDGFRA together with Total and phospho-AKT levels were analyzed by immunoblotting in K-Pα(+)S MSC and K-Pα(+)S KS cells after 72 hours of SAHA treatment. C) PDGFA and PDGFB levels were analyzed by immunoblotting in K-Pα(+)S MSC and K-Pα(+)S KS cells after 72 hours of SAHA treatment. Actin was used as loading control. D) K-Pα(+)S KS cells were treated with o.iuM of PDGFR tyrosine kinase inhibitor IV or left untreated, and incubated in an Incucyte Zoom (Essen Bioscience) to acquire green fluorescence images. The proliferation graph is shown the mean ± SE from three replicates for each condition. E) K-Pα(+)S KS cells were treated with SAHA alone or in combination with 0.1μm of PDGFR tyrosine kinase inhibitor IV, and incubated in an Incucyte Zoom (Essen Bioscience) to acquire green fluorescence images. The proliferation graph is shown the mean ± SE from three replicates for each condition. F) Immunofluorescence analysis of K167 expression (red) in K-Pα(+)S KS cells treated with SAHA alone or in combination with 0.1 μΜ of PDGFR tyrosine kinase inhibitor IV for 72 hours, nuclei were counterstained with DAPI (blue). Infected cells are GFP-positive. G) SA β-gal staining was performed after 72 hours of SAHA plus 0.1 μΜ of PDGFR tyrosine kinase inhibitor IV treatment of K-Pα(+)S KS cells. H) Total and phospho-PDGFRA, Cyclin Di, PDGFA and PDGFB levels were determined by immunoblotting in K-Pα(+)S KS cells treated with SAHA alone or in combination with 0.10M of PDGFR tyrosine kinase inhibitor IV for72 hours. Actin was used as loading control.

### PDGFRA signaling is a critical pathway to overcome KSHV-driven oncogene-induced senescence and for the proliferation of infected cells

To test whether or not the PDGFRA activation, which is prominent in induced K-Pα(+)S KS cells but not in K-Pα(+)S MSC cells, is necessary for enabling cell proliferation upon KSHV lytic reactivation, we inhibited this pathway using a highly selective PDGFR tyrosine kinase inhibitor (PDGFR Tyrosine Kinase Inhibitor IV, IC50: PDGFR-α: IC_50_= 45 nM; PDGFR-β: IC_50_ = 4.2 nM; C-Abl: IC_50_ = 22 nM; c-Src: IC_50_ = 185 nM; VEGFR: IC_50_ = 3.1 μΜ; bFGFR-i: IC_50_ = 45.8 μΜ; EGFR: IC_50_ = >100 μM). We found that in the presence of the PDGFR tyrosine kinase inhibitor K-Pα(+)S KS cells were not able to continue proliferating upon lytic reactivation (Figure 6E). Importantly, the same concentration of inhibitor was unable to halt proliferation of K-Pα(+)S KS cells that had not been lytically induced (Figure 6D). Moreover, upon lytic reactivation, inhibition of PDGFRA activation correlated with decreased levels of the proliferation marker Ki67 (Figure 6F) and with increased levels of senescence, SA-β-Gal staining (Figure 6G). Molecular analysis showed that the inhibition of PDGFRA signaling leads to a downregulation of Cyclin D1 and the PDGFRA ligands, PDGFA and PDGFB (Figure 6H). Taken together, our data indicate that activation of PDGFR signaling during KSHV lytic reactivation plays a role in allowing the proliferation of infected cells. These data, implicate this pathway as essential for overcoming KSHV-driven oncogene-induced senescence and for promoting KSHV-infected cell survival and proliferation, thus allowing KSHV tumorigenesis to progress.

### MSC culture conditions favor viral production in KSHV-infected human MSCs, while pro-angiogenic KS-like culture conditions are permissive for PDGFRA signaling-mediated proliferation of infected cells

To validate our results of KSHV infection with murine MSCs in a relevant human infection system, we studied the effect of different culture conditions in KSHV-infected human MSCs that were shown to be bona-fide human cell targets of KSHV infection [35–38]. Bone marrow-derived Human MSCs were infected with rKSHV219 (K-hMSC) and subsequently cultured in either MSC media (K-hMSC MSC) or in KS-like pro-angiogenic conditions (K-hMSC KS). We monitored the ratios of latent to lytic KSHV infection by GFP and RFP expression using fluorescence microscopy as well as by quantifying the percentage of GFP versus RFP-positive cells by Flow Cytometry analysis (Figure 7A-C). We observed that in contrast to mouse MSCs and regardless of the culture conditions, infected human MSCs showed both KSHV latently and lytically infected cells (Figure 7A). K-hMSC growing in KS-like media displayed lower percentages of RFP-positive cells than KSHV-infected hMSCs growing in MSC media (Figure 7B-C). Since hMSCs have been shown to be permissive for KSHV productive replication [35], we quantified the production of infectious virions by infecting HEK AD-293 cells. We found that K-hMSCs grown in MSC media produced more virions than those grown in KS-like conditions (20% vs 5% of infected HEK AD-293) (Figure 7D). Quantification of KSHV DNA content in the supernatants of the cultures showed that K-hMSC MSC contained more KSHV genomes/mL than K-hMSC KS, which correlated with the higher infectious virions content of the supernatants (Figure 7E). Paradoxically, the KSHV pattern and levels of viral gene expression in K-hMSCs MSCs and KS were quite similar regardless of the culture conditions (Figure 7F), suggesting that these culture conditions may be affecting the efficiency of virions production.

**Figure 7.**
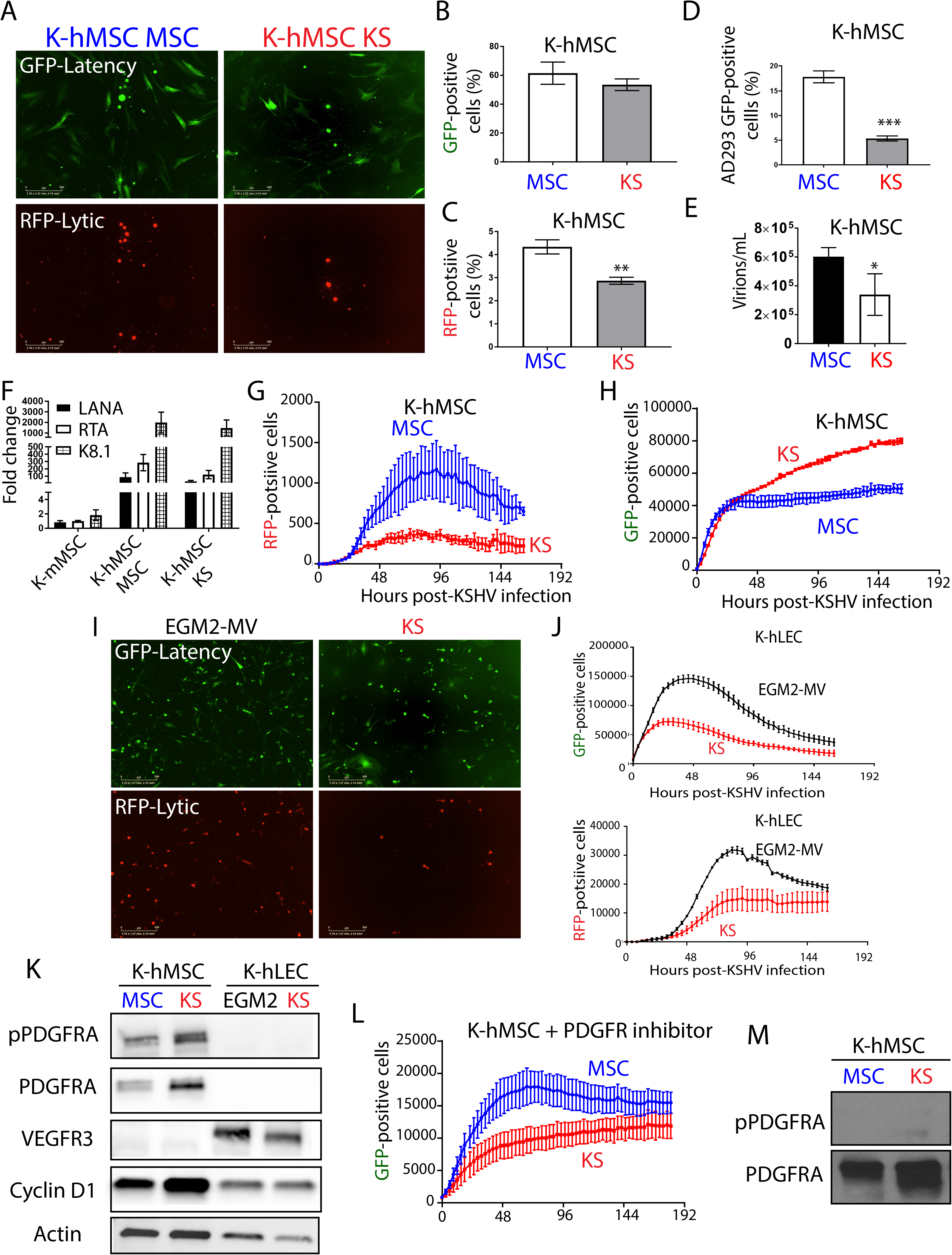
**MSC culture conditions favor viral production in KSHV-infected human MSCs, while KS-like culture conditions are permissive for PDGFRA-mediated proliferation of infected cells.** A) Human MSC KSHV infection was monitored by fluorescence microscopy using the GFP reporter driven by the constitutive promoter of cellular EF-1 and the RFP reporter driven by the early lytic gene PAN promoter. B) GFP expression analysis by flow cytometer in human MSCs after 96 hours of KSHV infection in MSC or KS-like media. The graph shown is from triplicates and presented as means ± SD. C) RFP expression analysis by flow cytometer on human MSCs after 96 hours of KSHV infection in MSC or KS-like media. The graph shown is from triplicates and presented as means ± SD. **P < 0.05. D) Cell-free supernatants of rKSHV.2ig-infected human MSC were used to *de novo* infect HEK AD293 cells. After 72 hours of infection, HEK AD293 GFP-positive cells were measured by flow Cytometry. The graph shown is from triplicates and presented as means ± SD. ***P < 0.05. E) Bars represent mean percent copy number of viral DNA levels in cell-free supernatants of human MSCs after 96 hours of KSHV infection in MSC or KS-like media determined by qPCR. The graph shown is from triplicates and presented as means ± SD. *P < 0.05. F) Fold-changes in KSHV gene expression between mouse K-Pα(+)S MSC cells, K-hMSC MSC and K-hMSC KS cells after 96 hours of KSHV *de novo* infection determined by RT-qPCR in triplicates and presented as means ± SD. G) Human MSCs were infected with KSHV in MSC or KS-like media and incubated in an Incucyte Zoom (Essen Bioscience), acquiring red fluorescence images. The proliferation of lytically infected cells (RFP-positive) is plotted overtime. The graph shows the mean ± SE from three replicates for each condition. H) Human MSCs were infected with KSHV in MSC or KS-like media and incubated in an Incucyte Zoom (Essen Bioscience), acquiring green fluorescence images. The proliferation of latently infected cells (GFP-positive) is plotted overtime. The graph shows the mean ± SE from three replicates for each condition. I) Human LECs KSHV infection in EGM2-MV or KS-like media was monitored by fluorescence microscopy using the GFP reporter driven by the constitutive promoter of cellular EF-1 and the RFP reporter driven by the early lytic gene PAN promoter. J) Human LECs were infected with KSHV in EGM2-MV or KS-like media and incubated in an Incucyte Zoom (Essen Bioscience), acquiring red fluorescence images, top panel, and green fluorescence images, bottom panel. The proliferation of lytically infected cells (RFP-positive, bottom panel) and latently infected cells (GFP-positive, top panel) is plotted over time. The graph shows the mean ± SE from three replicates for each condition. K) Total and phospho-PDGFRA, Cyclin D1 and VEGFR3 levels were determined by immunoblotting in human MSCs and LECs after 96 hours of KSHV infection in MSC, EGM2-MV or KS-like media. L) Human MSC cells were infected with KSHV in MSC or KS-like media and incubated in an Incucyte Zoom (Essen Bioscience) in the presence of o.i μm PDGFR tyrosine kinase inhibitor IV, acquiring green fluorescence images. KSHV infection and proliferation of the GFP positive cells is plotted over time. The graph shows the mean ± SE from three replicates for each condition. M) Total and phospho-PDGFRA levels were determined by immunoblotting in human MSCs after 96 hours of KSHV infection in MSC or KS-like media in the presence of 0.1 μm PDGFR tyrosine kinase inhibitor IV.

These results show that culture of K-hMSCs under different environmental conditions also can lead to different outcomes for KSHV lytic replication. Since in K-hMSC, KSHV productive lytic replication was spontaneous, we decided to use this opportunity to compare the impact of KSHV lytic gene expression on the proliferation of K-hMSCs growing in different culture conditions without having to resort to drug-based or HDACi lytic reactivation. We used an IncuCyte® Live-Cell Imaging and Analysis System to follow the growth of RFP-positive and GFP-positive populations of K-hMSCs growing in MSC or KS-like media. As observed for murine MSCs, the RFP-positive population of both K-hMSC MSC and K-hMSC KS increased in numbers and reached a plateau of cell proliferation (Figure 7G). Interestingly, as found in murine MSCs, the GFP-positive population of K-hMSC KS continue to proliferate over time (Figure 7H) while the GFP-positive population of K-hMSC MSC stopped proliferating shortly after KSHV infection (Figure 7H). We next sought to determine if proliferation upon infection in KS-like media was a unique feature of PDGFRA expressing KSHV targets such as hMSCs. We choose low passage primary human lymphatic endothelial cells (hLECs) which are PDGFR-negative and were shown to be permissive for KSHV latent and lytic infection [13, 62, 63]. Interestingly, LECs showed less lytic infection in KS media than in their specific cell culture media, EGM2-MV, (Figure 7I and 7J bottom panels). Importantly, there were not able to continue proliferating after KSHV infection in any of the two cell culture media (Figure 7J, top panel).

Seeking a molecular explanation for the unique ability of K-hMSC growing in KS-like conditions to proliferate, and based on our results depicted in Figures 5 and 6 using mouse MSCs, we evaluated cyclin D1 expression and PDGFRA activation in human lymphatic endothelial cells and human mesenchymal stem cells infected with KSHV and exposed to the different media. As shown in Figure 7K, proliferating K-hMSC growing in KS media showed much higher levels of PDGFRA activation and Cyclin D1 expression than K-hMSC growing in MSC media. As expected, LECs showed low levels of expression of Cyclin D1 and no expression of PDGFRA after KSHV infection, which correlated with less cell proliferation. We employed VEGFR3 as a specific lymphatic endothelial marker (Figure 7K). To determine whether the observed PDGFRA signaling activation was necessary for enabling cell proliferation of K-hMSC KS, we measured proliferation of K-hMSC MSC and K-hMSC KS in the presence or absence of a PDGFR tyrosine kinase inhibitor (PDGFR Tyrosine Kinase Inhibitor IV). We found that in both KSHV-infected hMSCs grown in presence of the inhibitor, PDGFRA signaling was abolished and the proliferation was inhibited shortly after infection (Figure 7L-M). This pattern of proliferation is reminiscent of K-hMSCs MSC, which were not substantially affected by the PDGFR inhibitor. These results illustrate that, as shown for mouse MSCs, the outcome for KSHV infection of human cells is strongly determined by the cell type and the culture conditions. While lymphatic endothelial cells, in EGM2-MV and KS media, and mesenchymal stem cells in MSC culture conditions favor viral production and do not proliferate, mesenchymal stem cells infected in KS-like culture conditions are more permissive for enabling PDGFRA-mediated proliferation of productively infected hMSC cultures.

## DISCUSSION

Human virally induced cancers are mainly characterized by long incubation periods and the fact that the majority of infected individuals do not develop cancer. This is due to the need for specific host conditions and factors that are necessary to enable a full viral-mediated transformation [1]. The KS spindle-cell progenitor and specific host conditions of KSHV-driven oncogenesis upon *de novo* infection are poorly understood. We identified PDGFRA(+)/SCA-1(+) bone marrow-derived mesenchymal stem cells (Pα(+)S MSCs) as KS spindle-cell progenitors and found that pro-angiogenic environmental conditions typical of KS, inflammation and wound healing are critical for KSHV sarcomagenesis. This is because growth in KS-like conditions generates a de-repressed KSHV epigenome allowing oncogenic KSHV gene expression in infected Pα(+)S MSCs. Furthermore, these growth conditions allow KSHV-infected Pα(+)S MSCs to overcome KSHV-driven oncogene-induced senescence and cell cycle arrest via a PDGFRA-signaling mechanism; thus identifying PDGFRA not only as a phenotypic determinant for KS-progenitors but also as a critical enabler for viral oncogenesis (Figure 8).

**Figure 8.**
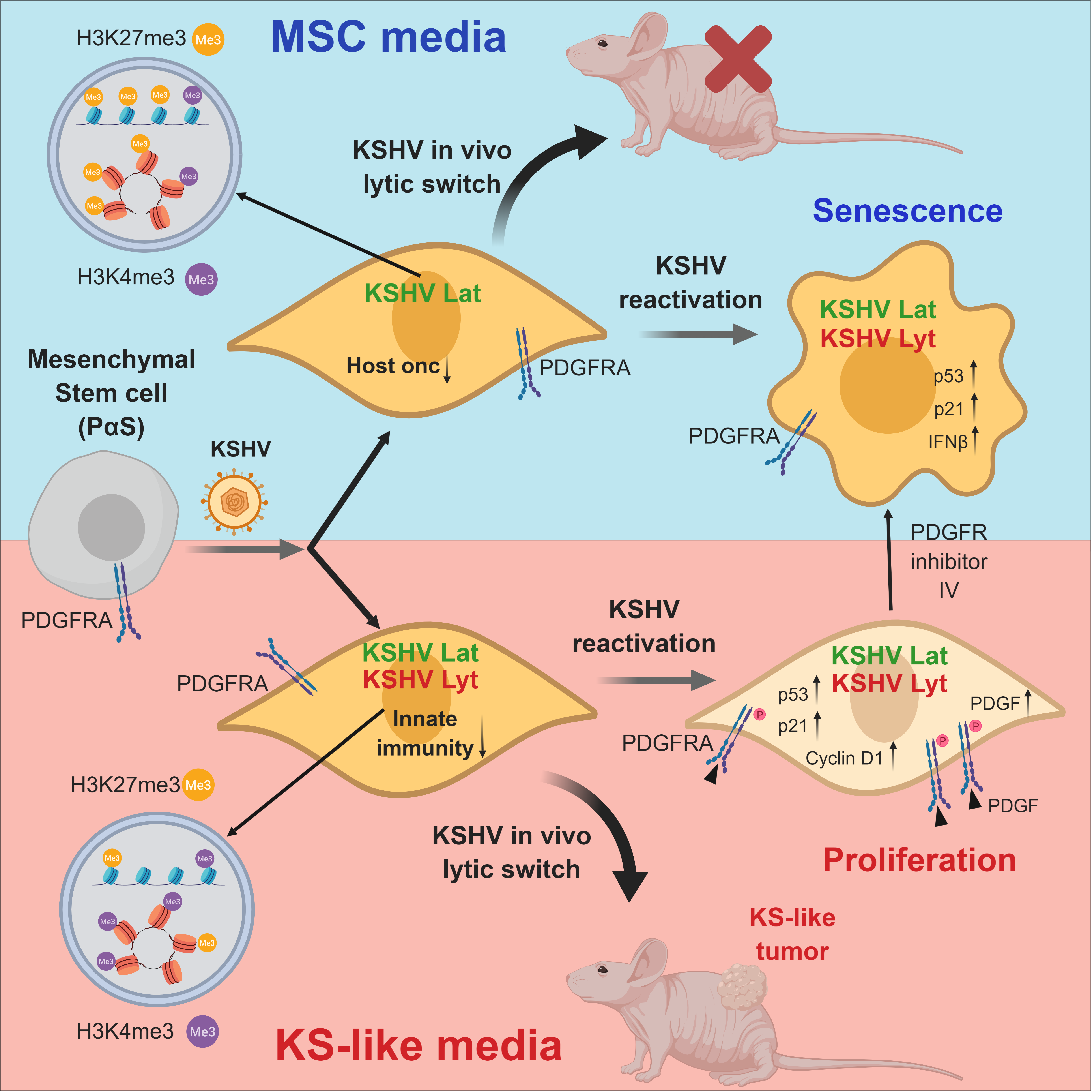
**PDGFRA Defines the Mesenchymal Stem Cell Kaposi’s Sarcoma Progenitors by Enabling KSHV Oncogenesis in an Angiogenic Environment.**Model showing PDGFRA(+)/SCA-i(+) bone marrow-derived mesenchymal stem cells (Pα(+)S MSCs) as KS spindle-cell progenitors. Pro-angiogenic environmental conditions typical of KS (KS-like media), inflammation and wound healing are critical for KSHV sarcomagenesis. This is because growth in KS-like conditions generates a de-repressed KSHV epigenome allowing oncogenic KSHV gene expression in infected Pα(+)S MSCs. Furthermore, these growth conditions allow KSHV-infected Pα(+)S MSCs to overcome KSHV-driven oncogene-induced senescence and cell cycle arrest via a PDGFRA-signaling mechanism; thus identifying PDGFRA not only as a phenotypic determinant for KS-progenitors but also as a critical enabler for viral oncogenesis.

AIDS-KS lesions are characterized by proliferating KSHV-infected spindle cells, however, the origin of the spindle cells (SCs) remains enigmatic because they express markers of multiple cellular lineages, including endothelial, monocytic, and smooth muscle [6], Initial immunohistochemistry studies showed that KS SCs are poorly differentiated cells showing phenotypic markers such as VEGFRs that suggest an endothelial origin, however, KSHV-infected primary endothelial cells in culture do not outgrow their uninfected counterparts, lose their KSHV genomes after extended passage in tissue culture, do not grow in soft agar and do not form tumors in nude mice [64]. On the other hand telomerase-immortalized human endothelial cell line infected with KSHV can become transformed and tumorigenic [24, 25]. Prior studies have suggested the role of mesenchymal stem cells (MSCs) as KSHV target and KS progenitors [28, 34–38]. In fact several KS models made from primary bone marrow-derived mouse endothelial lineage/adherent cells [26, 27] and one from rat MSCs [28] formed tumors in a KSHV-dependent manner, suggesting that these populations contain cell types in which KSHV infection is oncogenic. Sarcomas are cancers of mesenchymal origin [31], with Platelet-derived growth factor receptor (PDGFR) signaling playing a significant part in mesenchymal biology, including mesenchymal stem cell differentiation, growth, and angiogenesis [30]. Moreover, our recent studies showing that PDGFRA signaling is an oncogenic driver in KSHV sarcomagenesis and that it is consistently found in AIDS-KS tissue samples in its activated form [14], point to a bone marrow-derived PDGFRA(+) MSCs such as the Pα(+)S MSCs, which our study now show is a cell progenitor target of KSHV-driven sarcomagenesis.

Clinical observations, such as the appearance of KS tumors in surgical scars or sites of trauma *(Koebner* phenomenon)[65], suggest that cellular mechanisms such as inflammation, wound repair and angiogenesis are necessary to promote KS tumorigenesis. We found that Pα(+)S MSCs, which are recruited to sites of injury and inflammation [66], are highly permissive to KSHV infection. More importantly, we found that this infection is only oncogenic when it occurs in the context of a pro-angiogenic KS-like environment. These conditions favor enhanced KSHV lytic gene expression in KSHV-infected Pα(+)S MSCs conferring malignant cell characteristics and tumorigenicity in nude mice (Figure 8). We previously found that KSHVBac36 is angiogenic and tumorigenic in an endothelial cell lineage of adherent bone marrow cell preparations [26], which prompted us to ‘target’ putative progenitors among this transfected heterogeneous cell population. Our present study identified the KS progenitor cell type in the bone marrow as the Pα(+)S MSCs, and demonstrated the need for pro-angiogenic KS-like culture conditions for oncogenic viral gene expression and tumorigenesis (Figure 8). To reproduce pro-angiogenic KS-like culture conditions we added to the cell culture media crude preparations of endothelial cell growth factors (ECGF) [41, 42] together with heparin, which was shown to potentiates the mitogenic activity of ECGF [67]. The main active component of these crude extracts of ECGF is basic fibroblast growth factor (bFGF), which is a very strong mitogen for endothelial cells in culture and an angiogenic growth factor [68, 69], Several lines of evidence suggest that KS is an angiogenic and inflammatory cytokine-mediated driven disease; at least in early stages, and that angiogenic factors and; in particular, bFGF, play a role in lesion development [70, 71]. Basic FGF is highly expressed at the RNA and protein level by cultured AIDS-KS cells and in spindle cells from AIDS-KS and classic KS tissue [72, 73]· Moreover, *in vitro* studies have shown that a cooperation between PDGFB and bFGF is responsible of inducing the expression of PDGFRA and FGFR1/2, promoting proliferation of Pα(+)S MSCs without altering their multipotency or inducing their differentiation [74]. Our results point to the importance of these ECGFs as part of an angiogenic environment promoting enhanced oncogenic KSHV gene expression leading to KSHV oncogenesis.

Tumors that arise from K-Pα(+)S KS cells have an increased expression of KSHV oncogenes (KSHV *in vivo* lytic switch), paralleling our previous findings in the mouse KS-like mECK36 model [26], and also displayed PDGFRA signaling activation that co-distributed with KSHV LANA, thus supporting the described link between KSHV and PDGFRA activation in the tumors [14]. The pattern of KSHV expression in K-Pα(+)S KS cells-derived tumors show an increase in lytic transcripts *in vivo* [26]. The importance of the “in vivo” microenvironment on affecting the KSHV expression profile was previously reported in PEL and KS [24, 75]. Although it is assumed that most spindle cells in KS lesions are latently infected [76], KSHV lytic gene expression was also reported for a portion of KS lesions [43, 77, 78]. The RNA-sequencing analysis for KSHV gene expression of recently reported samples [43] shown in of Figure 2B, indicates that some human KS biopsies also express lytic transcripts as found in our mouse KSHV model, further supporting the relevance of KSHV *in vivo* lytic gene expression observed in MSCs tumors (the *in vivo* lytic switch) to actual cases of human AIDS-KS. We propose that the fact that tumorigenic K-Pα(+)S KS cells showed enhanced expression of KSHV lytic genes while they are able to growth *in vivo,* together with the results showing proliferation of productively KSHV-infected human MSC cultures of Figure 7, provides a powerful model for the initial steps of KSHV oncogenesis showing a definitive oncogenic role of KSHV lytic gene expression in KS pathogenesis.

The bioinformatics-based analysis using RNA sequencing of the mouse KS-like K-Pα(+)S tumors revealed glycolysis, NOTCH signaling and TGFB signaling among the most upregulated pathways during the KSHV *in vivo* lytic switch, pointing, therefore, to these pathways as important drivers of KSHV tumorigenesis *in vivo.* Indeed, many of these host signaling cascades that are co-opted by KSHV, including PI3K/AKT/mTOR,, NFkB and Notch, are critical for cell-specific mechanisms of transformation [1, 3, 79, 80]. Importantly, recent RNA sequencing data of KS transcriptomes compared to KS lesions show KSHV-mediated global transcriptional reprogramming that, similarly to our MSC tumors, included upregulation of the transforming growth factor-beta 1 (TGFB1) signaling pathway and glucose metabolism disorders [43].

The different KSHV transcriptional programs shown by K-Pα(+)S cells were determined by whether the culture conditions where the infected cells were grown was either those for MSC pluripotent growth or pro-angiogenic KS-like media. A similar pattern of differences in the KSHV transcriptional program was shown for KSHV-infected lymphatic endothelial cells (LECs) compared to KSHV-infected blood endothelial cells (BECs) [63]. Moreover, we showed that multiple biological processes were enriched among the differential expressed host genes (DEGs) between K-Pα(+)S MSC and K-Pα(+)S KS cells, indicating that KS-like culture conditions which favored the transcription of KSHV lytic genes correlated with the expression of oncogenic host-gene networks. Patterns of KSHV and host gene expression in K-Pα(+)S MSC versus K-Pα(+)S KS cells could be explained by differences in the epigenetic regulation determined by the different culture conditions. Here, we confirm that there were less repressive histone marks and more activating histone marks (H3K27me3-H3K4me3) on different portions of the KSHV genome in K-Pα(+)S KS than in K-Pα(+)S MSC cells, that correlated with increase KSHV gene expression by these tumorigenic cells (Figure 8). Pathway analysis of host genes enriched with H3K27me3 in non-tumorigenic K-Pα(+)S MSC showed that the most repressed genes were all related to KS and KSHV oncogenic signaling, further indicating that distinct environments where KSHV infection occurs can lead to different host epigenetic landscape, thus prompting oncogenesis. In the K-Pα(+)S KS cells, the most repressed pathways were linked to innate immunity, this could favor the ability of these cells to continue growing in the presence of increased levels of KSHV gene expression while promoting lower levels of expression of innate immune genes upon lytic reactivation. This indicates that tumorigenic K-Pα(+)S KS cells are more adapted to withstand oncogenic KSHV lytic gene expression possibly by upregulating oncogenic pathways, but also by repressing innate immune response genes upon induction of the lytic replication program (Figure 8).

A possible mechanism of oncogenicity in K-Pα(+)S KS cells is that these pro-angiogenic growth conditions promote a de-repressed KSHV epigenome, which during the *in vivo* KSHV lytic switch, allow for expression of oncogenic KSHV genes leading to tumorigenesis. We found that, as shown for other infected murine cells [81] neither viral DNA replication nor virus production was seen after KSHV reactivation with HDAC1 in K-Pα(+)S MSC and K-Pα(+)S KS cells, further indicating an abortive lytic state after reactivation characteristic of KSHV infection of murine cells. Moreover, K-Pα(+)S MSC cells stopped proliferating after KSHV lytic reactivation *in vitro* and *in vivo* in Matrigel-plugs. Interestingly, tumorigenic K-Pα(+)S KS were able to continue proliferating after KSHV lytic reactivation and after KSHV *in vivo* lytic switch in Matrigel-plugs, in spite of enhanced KSHV oncogene expression. Evidence has accumulated of senescence as a contributor of tumor suppression and as a host defense mechanism in limiting the effects of oncogenic viruses [56, 82]. We found increased senescence markers (SA-β-Gal positive cells, upregulation of SASP and senescence markers, increase p21 expression and decrease Ki67 expression) in non-proliferating K-Pα(+)S MSC after KSHV lytic reactivation, indicating that this inhibitory mechanism could explain the arrest in proliferation and the lack of tumor formation of these cells. In contrast, tumorigenic K-Pα(+)S KS cells are able to proliferate in spite of p21 upregulation and enhanced KSHV lytic gene expression (Figure 8). Early steps of KSHV tumorigenesis involve activation of the DNA damage checkpoints that lead to selective pressure for mutations which abrogate checkpoints, thus providing an advantage for cells with defective DNA damage response components to propagate [57]. Accordingly, tumorigenic K-Pα(+)S KS cells are able to proliferate in the presence of DNA damage (H2AX) after KSHV reactivation. Thus, these conditions not only conferred an epigenetic adaptation leading to increased KSHV gene expression but also to a mechanism, which upon *in vivo* lytic switch, would allow cells to overcome KSHV-driven oncogene-induced senescence and cell cycle arrest, prompting tumorigenesis (Figure 8).

We sought to understand the mechanisms that, after KSHV lytic reactivation, allow K-Pα(+)S KS cells to continue growing while induced senescence of K-Pα(+)S MSC cells. An RTK array showed that PDGFRA was the most activated kinase in K-Pα(+)S KS comparesd to K-Pα(+)S MSC. Furthermore, the fact that K-Pα(+)S MSC display a series of RTKs that are more activated than K-Pα(+)S KS (ErbB2, ErbB3), further indicates the unique role of PDGFRA activation compared to activation of other RTK cascades in promoting proliferation in the context of KSHV lytic gene expression. We have recently shown that KSHV usurps sarcomagenic PDGFRA signaling to drive KS through upregulation of PDGFs ligands by KSHV lytic genes [14]. After KSHV reactivation, only K-Pα(+)S KS cells showed PDGFRA activation that correlated with an upregulation of PDGFA and PDGFB expression. This KSHV-induced ligand-mediated activation of PDGFRA signaling may constitute an important mechanism mediating the adaptation of the K-Pα(+)S KS cells to continue proliferating despite KSHV lytic gene expression.

We found that K-Pα(+)S KS cells treated with PDGFR tyrosine kinase inhibitor lost their ability to continue proliferating upon KSHV lytic reactivation while showing increased levels of senescence markers. Thus, the PDGFRA signaling pathway is essential to promote KSHV-infected cell survival and proliferation allowing KSHV tumorigenesis to progress, further validating this oncogenic pathway as a therapeutic target for early stages of KS tumorigenesis (Figure 8). One approach that has been explored for the treatment of KSHV tumorigenesis is the stimulation of lytic reactivation in the presence of antiviral drugs. We have previously shown that this was effective in blocking PEL growth in mouse xenograft models [50] by the combination of proteasome plus HDAC inhibitors. Hereby we propose that the combination of HDAC plus PDGFR inhibitors should also constitute a plausible therapeutic approach.

It has been reported that human MSCs supported active viral lytic replication at the acute infection stage with low proliferation rates [35]. We observed that KSHV-infected bone marrow-derived human MSCs (K-hMSCs) grown in MSC media produced infectious virions, and in accordance, stop proliferating shortly after infection. However, when maintained in KS-like pro-angiogenic environment, these KSHV-infected hMSCs proliferated at high rates even though they showed active viral lytic replication, which correlated with higher levels of Cyclin D1 expression and PDGFRA activation. In addition, when K-hMSCs were grown in the presence of the PDGFR inhibitor IV to abrogate PDGFRA signaling, their proliferation was abolished shortly after KSHV infection. These results illustrate that, as shown for mouse MSCs, the outcome for KSHV infection of bone marrow-derived human MSCs is strongly determined by the culture conditions. While MSC culture conditions favor viral production, KS-like culture conditions are more permissive for enabling PDGFRA-mediated proliferation of productively infected hMSC cultures.

Several prior studies have pointed to the role of Mesenchymal Stem Cells (MSCs) and Lymphatic Endothelial Cells (LECs) as KSHV targets and KS progenitors [13, 17, 28, 34–38]. These studies propose that Kaposi sarcoma spindle cells can arise from either KSHV-infected MSCs through a mesenchymal to endothelial transition (MEndT) or from KSHV-infected LECs through an endothelial to mesenchymal transition (EndMT) [83]. We showed that after KSHV de novo infection of MSCs and LECs, all cells were lytically infected but only MSCs growing in KS-like environment were able to proliferate. The results of Figure 7 suggest a double role for these cells in KSHV pathobiology: pluripotent MSCs may play a role in supporting KSHV infection/ dissemination, while the same cells in a pro-angiogenic KS-like environment might be key precursors in KSHV sarcomagenesis. Regarding the proposed role of LECs as KS-progenitors; infected LECs have been shown to undergo EndMT when growing in “in vivo” -like conditions such as 3D spheroids and activation of notch signaling [13]. It then becomes plausible that an infected LEC could be also a KS progenitor through a first stage involving EndMT to become a mesenchymal PDGFR-positive KS-precursor that may undergo sarcomagenesis in an angiogenic KS-like environment. Our results point to PDGFRA as both the key pathogenesis driving receptor and the defining marker of the phenotype of an uninfected circulating KS progenitor (PDGFRA-positive bone marrow-derived MSCs), therefore intervening through the different steps of KSHV oncogenesis. PDGFs are the most powerful MSC chemo-attractant [84] and are expressed in KSHV-infected cells in KS lesions [14, 85, 86], We propose that a pro-angiogenic KS-like environment rich in secreted lytic infection-driven growth factors, like PDGF, can recruit circulating Pα(+)S-derived MSCs KS progenitors infected with KSHV and lead to tumorigenesis driven by KSHV oncogenic gene expression.

In summary, we designed a novel model of “de novo” KSHV oncogenesis based on infection of PDGFRA-positive mesenchymal stem cell progenitors cultured in pro-angiogenic/vasculogenic KS-like growth conditions. This system serves as a unique and robust platform to identify KSHV oncogenic-pathways and their relationship with cellular lineages and extracellular growth environments. Additionally, it can further be exploited to genetically dissect and elucidate viral, host-cell and environmental features of KS pathobiology.

## MATERIALS AND METHODS

### Cell Culture

Mouse mesenchymal stem cells (MSCs) were harvested and characterized as described previously [39]. MSCs were grown in 20% FBS (Atlanta Biologicals), 1% (vol/vol) penicillin and streptomycin, and αMEM (Invitrogen). MSCs were characterized by (i) adherence to plastic, (ii) negative for hematopoietic cell surface markers CD34 and CD45 and positive for CD105, CD90.2, CD73, and stem cell antigen SCA-1; and (iii) the ability to differentiate into adipocytes or osteoblast-like cells. Human MSCs were isolated as described previously [39]. Briefly, BM aspirates (25-50 mL) were purchased from AllCells (Emeryville, CA) under appropriate informed consent and institutional review board approval. The experiments were performed using MSCs at passages 8-15 for murine cells and passages 4-7 for human MSCs. Human lymphatic endothelial cells (LECs) were purchased from ScienceCell (catalog # 2500),used at passages 2-4, cultured in fibronectin-coated culture vessels (2μg/cm^2^) and grown in EGM2-MV medium from Lonza (catalog # CC-3202).

### Animal Studies

All mice were housed under pathogen-free conditions. Tumor studies were done in 4- to 6-week-old nude mice obtained from the National Cancer Institute. Tumors were generated by subcutaneous injection of MSC PDGFRA+ve KSHV KS cells (2 x 10^s^ cells) as previously described [26]. Tumor volumes were measured using a caliper every 2 days and calculated using the following formula: [length (mm) x width (mm)^2^x 0.52][87].

### Generation and maintenance of mouse PDGFRA-positive MSCs grown in MSC or KS-like media

The isolated mouse MSC cells were infected with rKSHV.219, MOI of 3, in the presence of polybrene (8 μg/ml) for 1 hour. 2 days later, puromycin was added to the culture to select and expand the infected cells. All murine cells were cultured in MEM alpha media: supplemented with 20% heat inactivated fetal bovine serum (FBS) (MSC media); or KS-like growth medium (KS media): DMEM supplemented with 30% FBS (Gemini Bioproducts), 0.2 mg/ml Endothelial Cell Growth Factor (ECGF) (Sigma-Aldrich) or (ReliaTech), 0.2 mg/ml Endothelial Cell Growth Supplement (ECGS) (Sigma-Aldrich), 1.2 mg/ml heparin (Sigma-Aldrich), insulin/transferrin/selenium (Invitrogen), 1% penicillin-streptomycin (Invitrogen), and BME vitamin (VWR Scientific).

## VIRUS PREPARATION AND INFECTION

iSLK-219 cells harboring recombinant KSHV 219 were used for virus preparation. Briefly, infectious viruses of the 219 strain were induced from the respective iSLK cells by treatment with doxycycline and sodium butyrate for 4 days. The culture supernatants were filtered through a 0.45-μιτι filter and centrifuged at 25,000 rpm for 2 h. The pellet was re­suspended in phosphate-buffered saline (PBS), aliquot, and stored at *−70°Q* as infectious KSHV preparations. Virus infection was performed according to the method used in a previous study, with minor modifications. Mouse MSCs, Human MSCs and human LECs were seeded at 6 x 10^4^ cells per well in 6-well culture plates. After a day of culture, the culture medium was then removed and cells were washed once with PBS. The prepared KSHV inoculum, MOI of 3 for mouse MSCs and MOI of 8 human MSCs and LECs, and 8 μg/ml of Polybrene were mixed and added to the cultured cells. After centrifugation at 700 x *g* for 60 minutes, the inoculum was removed after 3 hour and 2 ml of culture medium was added to each well. For titration of infectious virions, HEK AD-293 cells seeded in 12 wells for 24 hours were infected with 1 ml of supernatants. At 72 hours post-infection, the GFP-positive cells were counted by Flow Cytometry.

## DETECTION AND QUANTIFICATION OF VIRION DNA

After 96 hours, post-infection, viral loads (KSHV DNA copy numbers) were determined by real-time quantitative PCR of cell-free supernatants (virions). Cell-free supernatants were also collected and used for *de novo* infection of AD293 cells in the presence of 5 ug/ul Polybrene.

### Reagents

Antibodies: Histone H3K27AC (cat. #4353), Histone H3 (cat. #4499), AKT, and p-AKT (cat. #4060), Ki67 (cat. #9129) and H2AX (cat. #9718) were purchased from Cell Signaling Technology (Danvers, MA); p21 (cat. #ab109199) and LANA (cat. #ab4103) from Abeam (Cambridge, MA), PDGFA (cat. #sc-9974), PDGFB (cat. #sc-365805), Cyclin D1 (cat. #sc-718), FLT4/VEGFR3 (cat. #sc321) and P53 (cat. #sc-6243) from Santa Cruz Biotechnology; Actin (cat. #A5441) from Sigma; and p-PDGFRA (Y742) (cat. #AF2114) and total PDGFRA (cat. #AF-3O7) from R&D Systems (Minneapolis, MN). Anti-CD31 (PECAM1) (cat. #550274) from BD Biosciences. PDGFR tyrosine kinase inhibitor IV (cat. # sc-205794) was purchased from Santa Cruz Biotechnology.

### Matrigel-plug

We re-suspended 2 × 10^6^ of K-Pα(+)S MSC or K-Pα(+)S KS cells in 500 μL phenol red-free Matrigel (BD Bioscience, cat. # 356235). The mixture was implanted subcutaneously into the back of 7 weeks old male NUDE mice (n =3 in each group) following the BD Bioscience protocol. Two implants were injected per mouse. As a negative control, we injected only Matrigel (n = 3). After 1 week, the mice were euthanized, and the Matrigel plugs were removed.

### Whole-animal fluorescent imaging

Injected mice were analyzed with IVIS spectrum whole live-animal imaging system (Perkin Elmer Inc., Waltham, MA, USA). Mice were anesthetized with isoflurane using a vaporizer, and a fluorescence image was obtained using GFP filter set (excitation wavelength, 488 nm; emission wavelength, 510 nm).

### Flow Cytometry

For analysis of PDGFRA expression, cells were diluted in FACS buffer of PBS containing 2% FBS and 1 mM EDTA. APC-conjugated anti-PDGFRA antibody (eBioscience, cat. # 17-1401-81), PE-conjugated anti-Sca-1 antibody (eBioscience, cat. #12-5981-81), or the isotype control were diluted in FACS buffer at 1:200, and added to the cells for 30 min incubation at 4C. Cells were then washed twice in cold PBS and fixed with 4% paraformaldehyde. All flow cytometry analysis was performed on a Becton-Dickinson LSR analyzer (BDBiosciences) and analyzed using FlowJo (Tree Star, Inc.) software.

### RNA-Sequencing analysis

RNA was isolated and purified using the RNeasy mini kit (Qiagen). RNA concentration and integrity were measured on an Agilent 2100 Bioanalyzer (Agilent Technologies). Only RNA samples with RNA integrity values (RIN) over 8.0 were considered for subsequent analysis. mRNA from cell lines and tumor samples were processed for directional mRNA-seq library construction using the Preparation Kit according to the manufacturer’s protocol. We performed paired-end sequencing using an Illumina NextSeq500 platform. The short-read sequences were mapped to the mouse reference genome (GRCm38.82) by the splice junction aligner TopHat V2.1.0. We employed several R/Bioconductor packages to accurately calculate the gene expression abundance at the whole-genome level using the aligned records (BAM files) and to identify differentially expressed genes between cell lines and cell lines and tumors.

Briefly, the number of reads mapped to each gene based on the TxDb. Mmusculus gene ensembls were counted, reported and annotated using the Rsamtools, GenomicFeatures, GenomicAlignments packages. To identify differentially expressed genes between cell lines and tumor samples, we utilized the DESeq2 test based on the normalized number of counts mapped to each gene. Functional enrichment analyses were performed using the ClueGo Cytoscape’s plug-in (http://www.cytoscape.org/) and the InnateDB resource (http://www.innatedb.com/) based on the list of deregulated transcripts between K-Pα(+)S KS cells and K-Pα(+)S MSC cells (p-value<o.oi; FC>±i.s) and between K-Pα(+)S KS tumors *in vivo* and K-Pα(+)S KS cells *in vitro* (p-value <0.001; FC>±2). Data integration and visualization of differentially expressed transcripts were done with R/Bioconductor. KSHV transcriptome was analyzed using previous resources and KSHV 2.0 reference genome [88], while edgeR test was employed for differential gene expression analysis of KSHV transcripts. Kaposi’s sarcoma KSHV RNA-seq profiles were retrieved from GEO database (GSE100684) from a previous study [43] and integrated with mouse derived samplesforfurtheranalysis.

Functional enrichment analyses were performed using the ClueGo Cytoscape’s plug-in (http://www.cytoscape.org/) and the InnateDB resource (http://www.innatedb.com/) based on the list of deregulated transcripts between K-Pα(+)S KS cells and K-Pα(+)S MSC cells (p-valueco.oi; FC>±i.s) and between K-Pα(+)S KS tumors *in vivo* and K-Pα(+)S KS cells *in vitro* ip-value <0.001; FC>±2). Data integration and visualization of differentially expressed transcripts were done with R/Bioconductor.

### Chromatin immunoprecipitation (ChIP)

To reduce the effect of technical variation and sample processing bias, we utilized recently described quantitative ChIP (qCHIP) sequence technology. For normalization strategies, we obtained *Drosophila melanogaster* chromatin (Spike-in Chromatin) and Drosophila-specific histone variant, H2AV antibody (spike-in antibody) from Active Motif (cat. # 61686 and cat. # 53083). The ChIP protocol used previously was modified [48] (to perform the qChIP). In brief, for each ChIP, chromatin prepared from 10 million cells was mixed with 50 ng Spike-in chromatin. This was precleared using IgG for 2 hours before being mixed with 2 μg spike-in antibody and 4 μg antibody specific for the epitope of interest. The antibodies used were H3K4me3 (Abcam, cat. #ab8580), H3K27me2/3 mAb (Active Motif, cat. # 39535), control rabbit IgG (Santa Cruz Biotechnology, cat. #sc-2027). The antibody-chromatin mixture was incubated at 4C overnight in a cold room. Next day, 20 μ1 protein A/G magnetic beads (Pierce, cat. # 88802) were washed, blocked with PBS 1% BSA, added to the chromatin-antibody complex, and incubated at 4C for 2 hours. The supernatant was removed by placing the sample on a magnet. The beads were washed 4X with LiCl wash buffer and 1X with TE buffer. The DNA was eluted from the immune complex bound on beads by incubating the beads at 65C in freshly prepared 100 μ1 SDS elution buffer for two hours. Another elution step was performed by incubating the beads in 50 μl elution buffer. The two portions of eluted DNA were combined (150 μl) and DNA was finally purified using NEB Monarch PCR and DNA cleanup kit (cat. #T1030S). The DNA was finally eluted in 60 μl o.iX TE. 1 μΙ of eluted DNA was utilized for ChIP-seq library preparation.

### ChIP-seq library preparation

The NEB Next Ultra II DNA Library Prep Kit for illumina (cat. # E7645L) was used for preparing libraries with the manufacturer’s protocol and associated reagents. 50 μl eluted ChIP DNA for each sample was mixed with 3 μl of end prep enzyme mix and 7 μl end prep reaction buffer. The end prep was carried out by incubating the reaction for 30 min at 20C and 30 min at 65C in a thermocycler. The above reaction was mixed with 2.5 μl of 1:25 dilution of lllumina adapter, and ligated using 60 μl ligation master mix and ligation enhancer by incubating the reaction at 20C for 15 min. 3 μl of USER enzyme was added to carry out the reaction at 37C for 15 min. Libraries were purified to remove unligated adaptors using AMPure XP beads. The PCR amplification of purified adaptor-ligated fragments was performed using Q5 DNA Master mix supplied in the kit. In total 16 libraries were generated and were multiplexed using NEB Multiplex Oligo sets for Illumina. The libraries were pooled in two lanes of lllumina and were sequenced on lllumina HiSeq using 2*150 run at the University of Florida ICBR facility.

### KSHV ChIP-Seq analysis

The adapter sequences were trimmed from the reads using Trimmomatic [89] and the quality of the trimmed reads was checked with FastQC. Reads were aligned to the KSHV genome (GenBank accession number: GQ994935.1) [90] using Bowtie2 in paired-end mode [gi], Duplicates were removed with Picard, after which peaks were called using MACS2 [92].

The peaks were scaled with respect to KSHV episome copy number and *Drosophila* (fly) spike-in reads. To do the normalization, first, the data was normalized to the number of episomes.

To obtain the scaling factor, the number of episomes in the MSC input was divided by the number of episomes in the KS input and the MSC input. This gave the KS samples a scaling factor of approx. 1.8 and the MEM samples a scaling factor of 1. Next, the fly scaling factor was determined. This was done with the following equation for all samples: 1/(number of fly reads/1000000). The ratio of KSHV reads to fly reads was determined by dividing KSHV reads by fly reads. Much like step one, the episome and fly scaling factor was determined by dividing the MSC input KSHV/fly ratio by the KS and MSC KSHV/fly ratios. Finally, the spike in episome count was determined by multiplying the fly scaling factor and the KSHV/fly ratio. The data were visualized using IGV.

To generate the heat maps, promoter regions were defined as 1 kb upstream and downstream of the translational start sites (TSS). The TSSs were determined using ORF start positions and annotatePeaks.pl from Homer (V4.7), which resulted in an expression matrix. The rows display the histone modification patterns along the −1 kb to +1 kb genomic regions relative to the translational start site (TSS) of each viral gene, which we assigned as the gene regulatory regions. The 1 kb regions were divided into twenty 50 bp fragments and genes were separated into IE, E, L, and latent gene classes. The resulting matrix was log transformed and row centered before using Pearson correlation and pairwise complete-linkage hierarchical clustering with Cluster 3.0. Clustered matrices were then visualized using Java TreeView (version 1.1.6r4) where rows show the log2 average peak derived from the average of the biological replicates of the ChIP-Seq experiments. Blue and yellow colors represent the lower-than-average and higher-than-average expression, respectively.

### Host gene ChIP-Seq analysis

ChIP-seq reads were analyzed following the AQUAS ChIP-seq bioinformatics pipeline using default parameters (Kundaje lab, https://github.com/kundajelab/chipsegpipeline). Briefly, FASTQ reads were aligned to the mouse mm9 genome using BWA vo.7.13 and duplicate reads were removed using Picard tools v1.126. Peaks and signal tracks were generated using MACS2.1 and filtered for blacklist regions identified by ENCODE. Bedtools V2.26.0 intersect was used to determine peak overlaps with RefSeq genes +2.5kb upstream of TSS regions to identify H3K27me3 and H3K4me3 enriched genes. NGS Plot V2.61 was used to generate density plots.

### Soft agar assay

Base agar was made by combining melted 1% agar with KS-like medium or MSC medium to give a 0.5% Agar/1X KS-like or MSC medium solution. 1.5 mL was added to each well of a 6 well plate and allowed to set. Five thousand cells were plated on top of base agar in 0.7% agar/2X KS-like or MSC medium in triplicate in 6-well plates. The cells were fed every 3 days with 1 mL of Ks-like or MSC medium. Colonies were photographed at 4 weeks. Only colonies larger than the mean size of the background colonies in the negative control wells were considered.

### Real-Time Quantitative PCR (RT-qPCR)

RNA was isolated with RNeasy Plus Kit (QIAGEN, Valencia, CA) with on columns DNase treatment. 500 ng of RNA was transcribed into cDNA using Reverse Transcription System (Promega, Madison, WI) according to the manufacturer’s instructions. RT-qPCR was performed using an ABI Prism 7000 Sequence Detection System (Applied Biosystems) with SybrGreen PCR Master Mix (Quanta Biosciences). In every run, melting curve analysis was performed to verify the specificity of products as well as water and -RT controls. Data were analyzed using the ΔΔCT method as previously described [26]. Target gene expression was normalized to GAPDH by taking the difference between CT values for target genes and GAPDH (ΔCT value). These values were then calibrated to the control sample to give the ΔΔCT value. The fold target gene expression is given by the formula: 2^−ΔΔCT^.

### Cell proliferation assay (IncuCyte)

Cells were plated in 6-well plates at 60,000 cells/well in 3 replicates. Cells were incubated in an Incucyte Zoom (Essen Bioscience), acquiring green and red fluorescence images at 10× every 2 hours. The Incucyte Zoom software was used to analyze and graph the results.

### Immunofluorescence Staining

Immunofluorescence assay (IFA) was performed as previously described [26]. Briefly, paraffin-embedded tissue sections were deparaffinized and rehydrated following antigen retrieval treatment. Cells were fixed in 4% paraformaldehyde for 10 min and washed with PBS. Tumor section and cells were permeabilized in 0.2% Triton-X/PBS for 20 min at 4°C. After blocking with 3% of BSA in PBS and 0.1% Tween 20 for 60 min, samples were incubated with Primary antibodies overnight at 4C. After PBS washing, samples were incubated with fluorescent secondary antibodies for 1 hour (Molecular Probes), washed and mounted with ProLong Gold antifade reagent with DAPI (Molecular Probes). Images were taken using a Zeiss ApoTome Axiovert 200M microscope.

### Western Blotting

Protein concentrations in cell and tumor lysates were quantified using the DC Protein Assay (Bio-Rad,). 20 μg of proteins were mixed with Laemmli buffer, boiled for 5 min, resolved by SDS-PAGE and transferred to PVDF membranes (Bio-Rad Laboratories). Membranes were blocked with 3% BSA for 1 hour and incubated with primary antibodies (4C, 16 hours). After 3 TBS/T washes, membranes were incubated with HRP-labeled secondary antibodies (Promega) for 1 hour at room temperature. Protein bands were developed using ECL Plus Detection Reagents (GE Healthcare). To analyze multiple proteins on the same membrane, membranes were washed with Restore PLUS Western Blot Stripping Buffer (Thermo Scientific) according to the manufacturer’s protocol.

### Phospho-Receptor Tyrosine Kinase (RTK) Array

R&D Systems’ Mouse Phospho-Receptor Tyrosine Kinase (RTK) Array Kit (Catalog # ARY014) was used to detect levels of phosphorylation of 39 RTKs in K-Pα(+)S MSC or K-Pα(+)S KS cells.

## QUANTIFICATION AND STATISTICAL ANALYSIS

### Statistical Analysis

Statistical significance of the data was determined using two-tailed Student’s t-test and 2way ANOVA for multiple comparisons. A p-value lower than 0.05 was considered significant. Statistical analysis was performed using GraphPad Prism 7. All values were expressed as means ± standard deviation.

### Ethics Statement

The animal experiments have been performed under UM IACUC approval number 16-093. The University of Miami has an Animal Welfare Assurance on file with the Office of Laboratory Animal Welfare (OLAW), National Institutes of Health. Additionally, UM is registered with USDA APHIS. The Council on Accreditation of the Association for Assessment and Accreditation of Laboratory Animal Care (AAALAC International) has continued the University of Miami’s full accreditation.

## ACKNOWLEDGMENTS

We would like to thank, Darlah Lopez Rodriguez and Vytas Dargis-Robinson for their technical support. The Oncogenomics Core Facility at the Sylvester Comprehensive Cancer Center for performing high-throughput sequencing and the Flow Cytometry Core Facility for assistance with Flow Cytometry and cell sorting analysis. Lucas J. Boatwright and UF ICBR for sequencing and bioinformatics support. We would like to thank Dr. Priyamvada Rai, from the department of Medicine at the University of Miami for usefull advice.

## Supporting information captions

S1 Figure. Tumorigenic analysis **of KSHV-infected and uninfected MSCs growing in MSC media.**

S2 Figure. Percentage of **KSHV de novo infection in PDGFRA-negative and PDGFRA-positive cells in MSC and KS condition.**

S3 Figure. Viral DNA **copy number in K-Pα(+)S MSC cells, K-Pα(+)S KS cells and tumors.**

