## Supplementary figures and images for "PDGFRA Defines the Mesenchymal Stem Cell Kaposi’s Sarcoma Progenitors by Enabling KSHV Oncogenesis in an Angiogenic Environment"

### S1 Figure

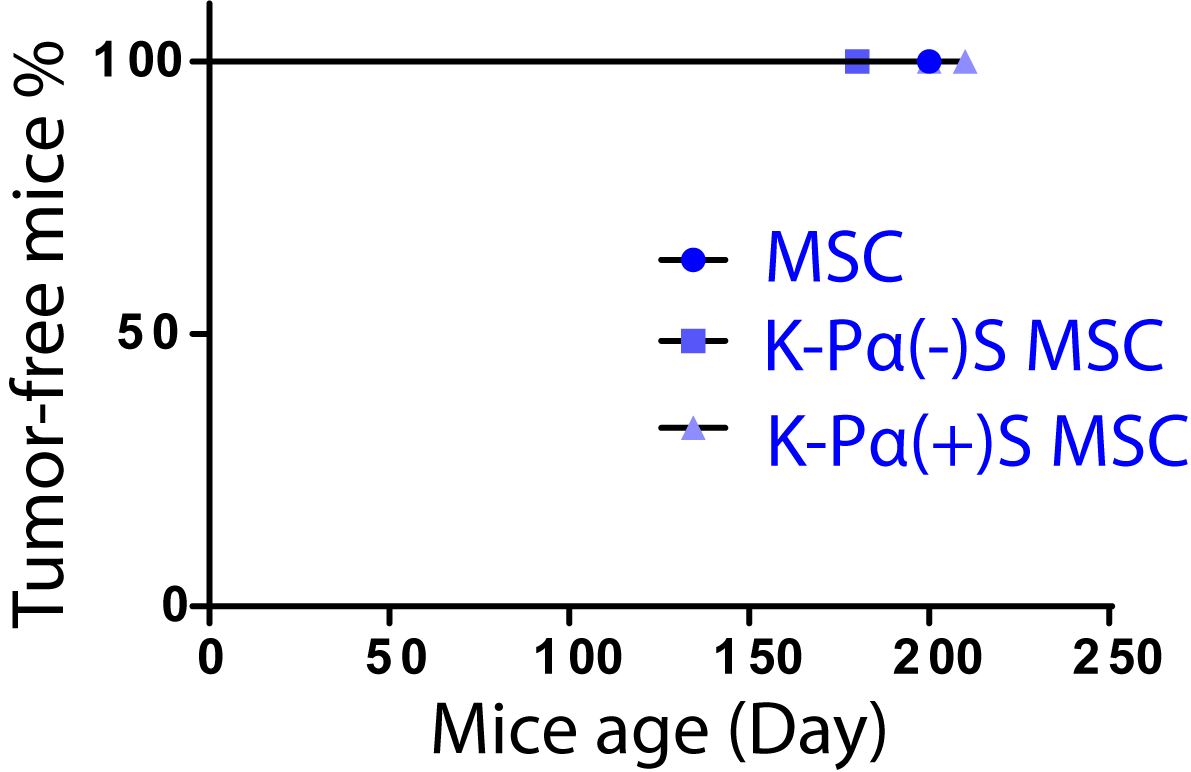

### S2 Figure

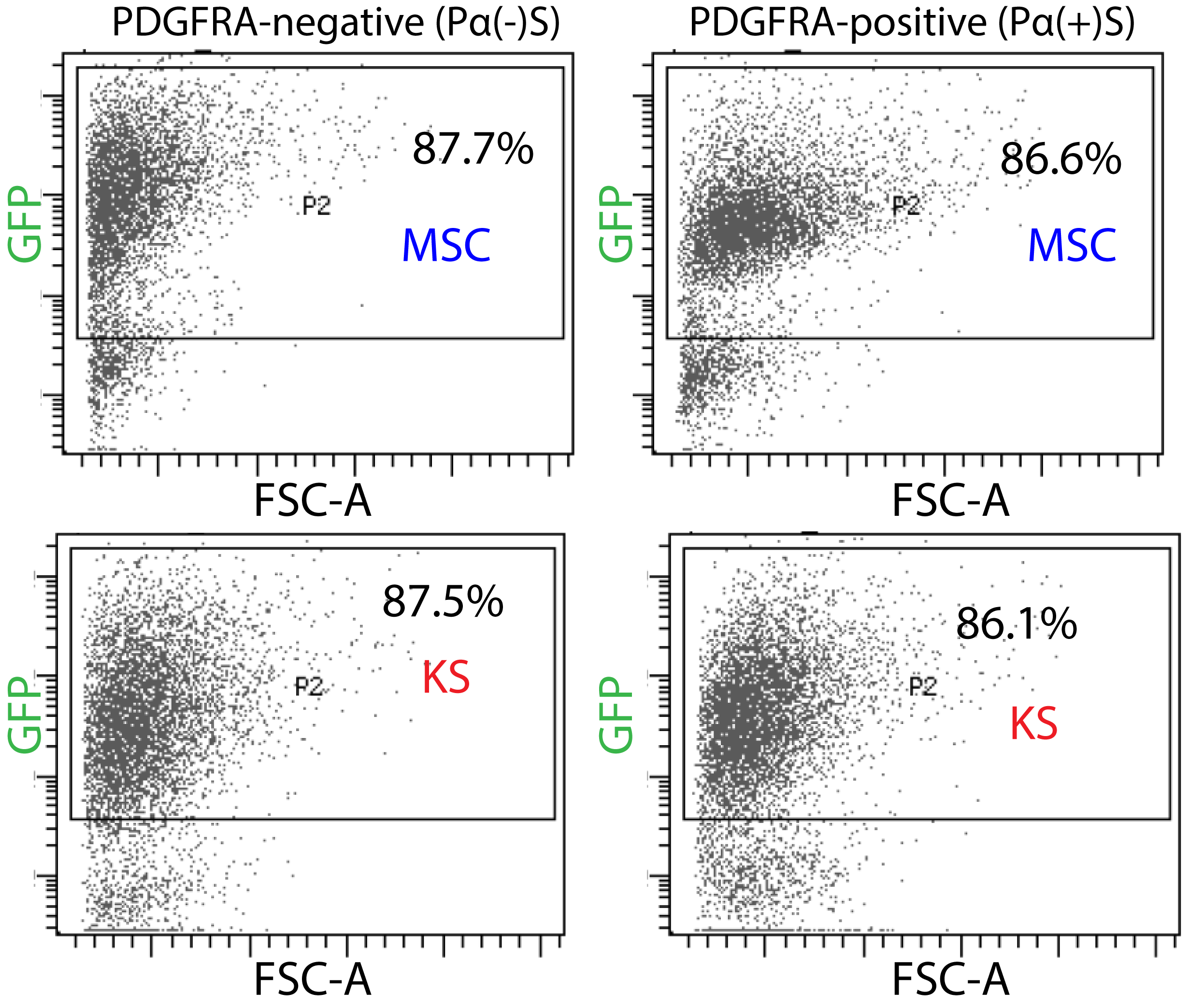

### S3 Figure

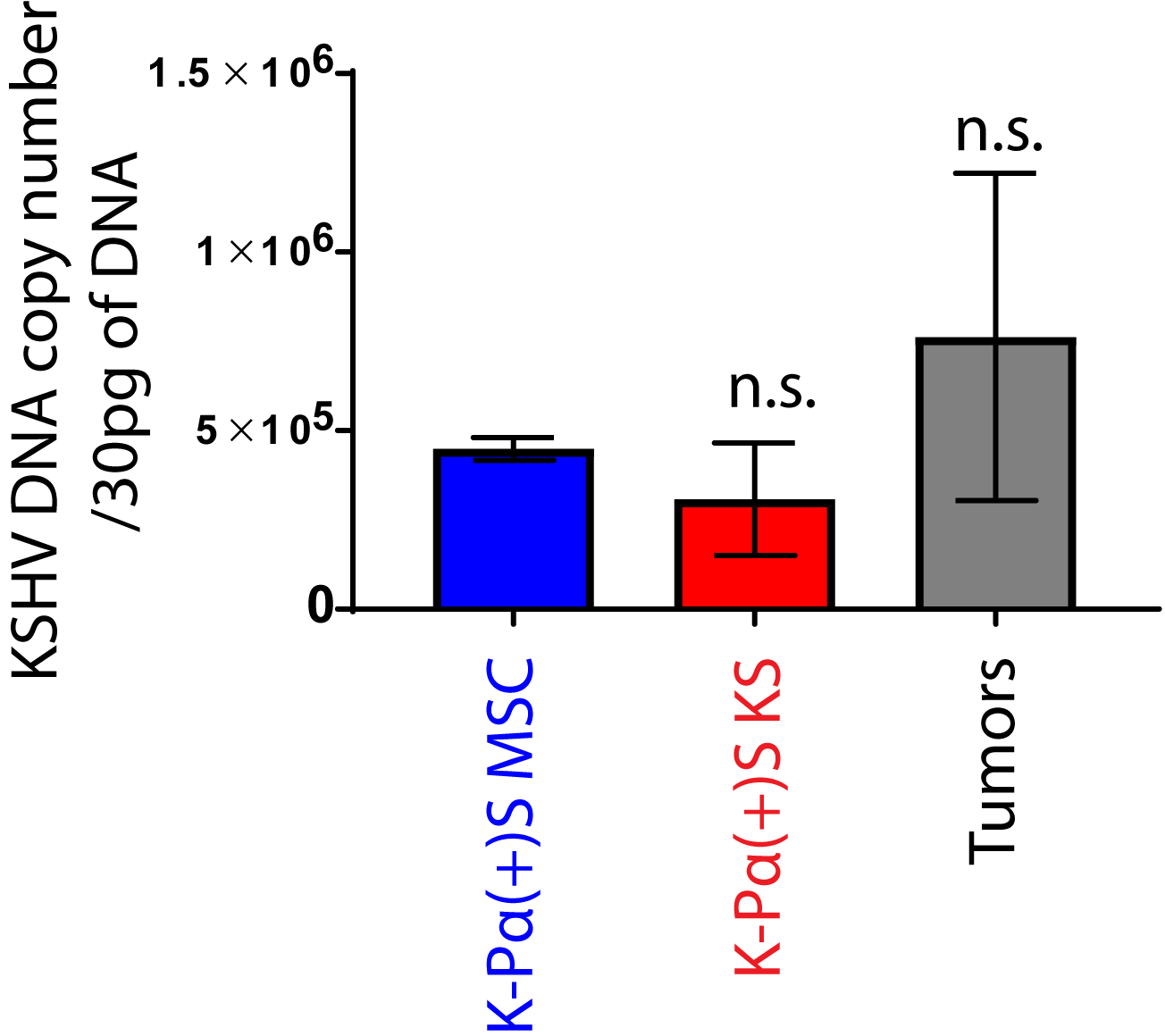
